# Foci, waves, excitability : self-organization of phase waves in a model of asymmetrically coupled embryonic oscillators

**DOI:** 10.1101/2024.06.24.600484

**Authors:** Kaushik Roy, Paul François

## Abstract

The ‘segmentation clock’ is an emergent embryonic oscillator that controls the periodic formation of vertebrae precursors (or somites). It relies on the self-organization at the Pre Somitic Mesoderm (PSM) level of multiple coupled cellular oscillators. Dissociation-reaggregation experiments have further revealed that ensembles made of such cellular oscillators self-organize into an oscillatory bidimensional system, showing concentric waves around multiple foci. Here, we systematically study the dynamics of a two dimensional lattice of phase oscillators locally coupled to their nearest neighbors through a biharmonic coupling function, of the form sin *θ* + Λ sin^2^ *θ*. This coupling was inferred from the Phase Response Curve (PRC) of entrainment experiments on cell cultures, leading to the formulation of a minimal Elliptic Radial Isochron Cycle (ERIC) phase model. We show that such ERIC-based coupling parsimoniously explains the emergence of self-organized concentric phase wave patterns around multiple foci, for a range of weak couplings and wide distributions of initial random phases, closely mimicking experimental conditions. We further study extended modalities of this problem to derive an atlas of possible behaviours. In particular, we predict the dominant observation of spirals over target wave patterns for initial phase distributions wider than approximately *π*. Since PSM cells further display properties of an excitable system, we also introduce excitability into our simple model, and show that it also supports the observation of concentric phase waves for the conditions of the experiment. Our work suggests important modifications that can be made to the simple phase model with Kuramoto coupling, that can provide further layers of complexity and can aid in the explanation of the spatial aspects of self-organization in the segmentation clock.

## I. INTRODUCTION

The study of self-sustained nonlinear oscillators and the exploration of their coupled collective behavior, such as synchronization, have long fascinated researchers from diverse disciplines, owing to their ubiquitous existence in a myriad of natural and engineered systems [1–6]. Among the tools from dynamical systems theory that have been used to understand their behavior, phase reduction theory [1, 7–10] has emerged as a pioneering mathematical framework that has allowed to simplify the analysis of complex oscillatory systems. This theory focuses on reducing the multidimensional dynamics of coupled nonlinear oscillators to a single, low-dimensional phase variable. In this series of papers, we aim to explore the dynamics of such phase oscillators which are coupled by a biharmonic coupling function of the form:

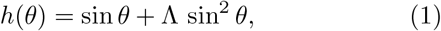

where *θ* represents the phase difference between two oscillators. As shown in Fig. 1, the sin^2^ *θ* term breaks the odd symmetry of the sin *θ* coupling of the original Kuramoto model [1, 9]. Another important thing to notice is that the coupling function stays mostly positive within the range of phases as we increase the degree of asymmetry, Λ. In this paper, we will be interested in the case when the oscillators are coupled only locally to their nearest neighbors in two dimensions. The results for the case of global all-to-all coupling will be presented in a separate paper.

**FIG. 1:**
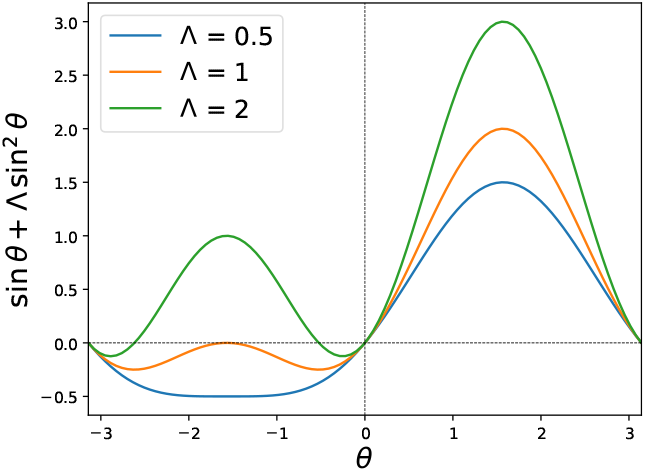
Behavior of the coupling function: *h*(*θ*) = sin *θ* + Λ sin^2^ *θ* for different Λ.

Our interest in studying this biharmonic coupling function has its roots in cellular oscillators implicated in embryonic development. The vertebrate segmentation clock [11, 12] is the global oscillatory gene expression controlling the sequential segmentation of the embryonic body axis into blocks of tissue called somites, which are vertebrae precursors (the overall process is called somitogenesis; see [13] for a recent review of existing models). The somitogenesis period or the rhythm of somite formation is fixed but species dependent e.g. 30 min in zebrafish [14], 90 min in chicken [11], 2.5 hr in mouse [15] and 5 hr in humans [16]. Zooming in, somitogenesis relies on coupled cellular oscillators in the presomitic mesoderm (PSM). Individual cells express a network of genes that interacts with a determination front [17–20] positioned by morphogen gradients. Multiple genes oscillate in a single cell oscillator, in particular in the Notch signalling pathway, which has been known to synchronize phases between neighboring PSM cells [21–27].

In recent years, following the pioneering work of [28], multiple groups have designed and studied so-called *exvivo* systems partially or fully mimicking somitogenesis in tissue culture. In Ref. [29], Sanchez *et al*. used a dynamical systems-inspired approach to study the entrainment dynamics and phase response of the segmentation clock, when subjected to weak external perturbations in the form of periodic pulses of DAPT [30]. DAPT is an inhibitor of the Notch signaling pathway. Using a dynamic fluorescent reporter for segmentation clock genes called LuVeLu [20], the effects on collective oscillations were observed using time-lapse microscopy with the intensity of the signal acting as a proxy for the phase of the clock. To quantify the influence of DAPT, a phase oscillator model was used. Phase oscillator models, wherein PSM cells are idealized as limit cycle oscillators with a single phase variable provide an expedient approach to investigate systems-level phenomena [28, 29, 31–39], including local and global synchronization. This method circumvents the requirement to introduce molecular (genetic) complexity into the problem, allowing for a focused exploration of the desired properties.

An asymmetric Phase Response Curve (PRC) was observed, meaning that oscillators qualitatively behave in the same way (delayed for DAPT) irrespective of the phase of their cycle. This is not expected from classical theory of oscillators close to Hopf bifurcations, where external perturbations applied at different phases of the cycle can either advance or delay the clock, leading to the classical Kuramoto coupling. Motivated by these insights, a model for the phase response, called the Elliptic Radial Isochron Cycle (ERIC) model was proposed in Ref. [29], whose infinitesimal PRC is precisely of the form given by Eq. 1. Importantly, it was proposed that such coupling reflects the properties of an internal oscillator poised close to a Saddle-Node on Invariant Cycle (SNIC) (or Saddle-Node Infinite Period (SNIPER)) bifurcation [29, 38]. Such asymmetric nature of the clock response is also consistent with the experimentally observed unidirectional nature of the Notch-Delta coupling, termed as the “walkie-talkie” model [40, 41]. This means that neighboring phases mostly “talk” to each other during a certain portion of their cycle, for example, when their relative phase difference is positive. Such asymmetric coupling has further been confirmed in the context of segmentation [42, 43].

Assuming we know both response and coupling properties of individual oscillators, one should be able to predict the emerging behavior of coupled oscillators. Two other experimental studies [44, 45] highlighted the self-organization capabilities of PSM cells to form spatiotemporal phase patterns. In Ref. [44], the entire PSM including both posterior and anterior regions of several mouse embryos were dissected, then dissociated into single cells and a randomized cell suspension was created where the PSM cells lost all positional information. When these dissociated cells were plated on fibronectin-coated coverglass and allowed to reaggregate, they formed multiple foci at characteristic distances from one another despite the randomization. Each such foci, termed emergent PSM (ePSM), resembled monolayer explants [28] with the center and the periphery resembling the posterior and anterior PSM respectively. Notably, each foci displayed concentric phase wave patterns with the oscillation dynamics dependent on the phase and frequency of the input population, rather than exhibiting special, segregated pacemaker cell populations. Similarly, the authors in Ref. [45] established novel culture conditions in which posterior mouse PSM explants displayed stable oscillations. This further enabled them to culture dissociated cells and study cell autonomous oscillatory properties which is an important assumption in the phase oscillator representation of PSM cells. Excitability properties were observed, controlled by the YAP signalling pathway [46–51], which is regulated by cell density. Once the threshold is exceeded, juxtacrine Notch signaling acts as the stimulus triggering the oscillations. In combination, these studies [44, 45] brought about important breakthroughs in our understanding of the segmentation clock properties both at the level of isolated cells and collective oscillations.

In this work, we will explore the phase dynamics of a population of ERIC model oscillators, which are coupled locally to their nearest neighbors in 2D. Assuming that the PSM cells (phase oscillators) in mouse explants and reaggregated populations of dissociated cells are coupled in this fashion, we can *parsimoniously and uniquely* recapitulate the collective self-organization dynamics observed in experiments involving mouse explant systems, thus bridging the response properties of single cells (Eq. 1 derived from at least three independent sets of experiments [29, 42, 43]) to the emergent properties of a cell population (observed in two other independent experiments[44, 45]).

The rest of the paper is organized as follows. In Sec. II, we define the dynamical equations for the locally coupled phase model using the asymmetric, biharmonic form of coupling, that was inferred from the entrainment experiment [29] on mouse PSM explants. We perform a comprehensive study of the dependence of the phase landscape on the initial conditions. The origin of these phase wave patterns are explored briefly in Sec. III using the discrete model and its continuum version. In Sec. IV, we introduce excitability into our phase model and connect the model parameters to the biological controls. For both models, we make theoretical predictions for the observation of spiral phase wave patterns and propose experiments that can be designed to observe them. Beyond the phasemaps, we have also explored other properties of the excitable ERIC model such as the coherence behavior at the local and global level. These results are presented in Sec. IV B. We conclude in Sec. V with a summary and discussion of the main results and provide an outlook on future work that can be done to provide extra layers of complexity in the modeling. We also provide an extensive supplement [52] that includes a detailed comparison to other phase models which fail to generate the phase patterns observed experimentally, details and hyperlinks to the Python codes necessary to generate the plots and animations presented in this work.

## II. DYNAMICAL EQUATIONS AND PHASE MAPS

### A. Setting up the model

We study the phase dynamics of a population of locally coupled ERIC model phase oscillators arranged in a square packing on a two-dimensional *N* × *N* lattice. Their intrinsic time-independent frequencies are represented by *ω*_*i*,*j*_ (*i, j* = 1, …, *N*) and we assume that nearest neighbors are all coupled to one another with identical strength *K* in the first harmonic and *K*Λ in the second harmonic. Λ is a dimensionless parameter that determines the relative strength of the second harmonic as well as the asymmetry in the coupling function (see Fig. 1). The phase dynamics of the oscillators in this locally coupled 2D ERIC model (2*DE*) are governed by the following equations,

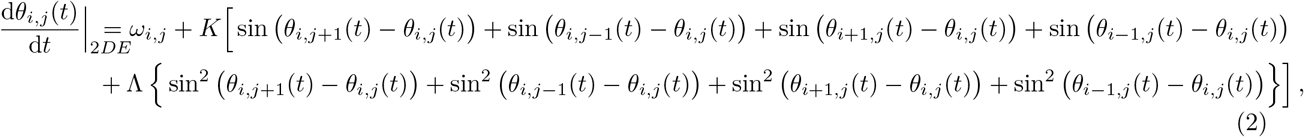

where *i, j* = 1, …, *N*. Here *θ*_*i*,*j*_(*t*) denotes the phase of the oscillator at the (*i, j*)-th lattice site at time *t*. This is an interesting problem to study in its own right because of the addition of the genuine higher-order harmonic in the coupling function. Here, we will instead focus on modeling qualitatively the spatiotemporal phase dynamics of mouse PSM cells observed in Refs. [44, 45]. The phase oscillators in the lattice are a coarse-grained representation of the genetic oscillators in the PSM cells. Below, we will state our rationale for choosing the initial conditions for simulating the dynamics.

In the randomization-dissociation experiments of Ref. [44], the whole PSMs of several mouse embryos are dissociated into single cells and the randomized cell suspension is allowed to form dense cell re-aggregates. These are then cultured on fibronectin-coated coverglass, and real-time imaging is used to quantify the signaling activity using the Notch reporter LuVeLu. We are exploiting this 2D geometry with the assumption that the oscillatory property of the PSM cells is intrinsic and remains independent of the spatial dimensionality. Apart from the randomness in initial conditions of the experiment, we take into account the experimentally well-established fact that there is a spatial period gradient along the embryonic axis, such that cells close to the posterior of the embryo oscillate faster (higher frequency) compared to those in cells located more anteriorly. This has been observed across species and also in *in vitro* assays that culture intact or dissociated PSM [28, 32, 35, 44, 53–55]. Also, the oscillation period of PSM cells in isolation is usually longer (for posterior mouse PSM cells, *T* ∼ 155 min) than what is observed *in vivo*, where they are coupled to other cells [24].

Based on these experimental observations, we will randomly sample the natural frequencies *ω*_*i*,*j*_ (*i, j* = 1, …, *N*) of the phase oscillators from a uniform distribution between [2*π/*180, 2*π/*150] min^−1^. This allows for the randomization performed in the experiment and also respects the period gradient arising from sampling cells across the PSM and from multiple mouse embryos. For the discussion below, we will consider a 50×50 square lattice corresponding to 2500 PSM cells and use open boundary conditions on the lattice for the phases [56]. In the supplementary material [52], we have provided results obtained for other choices of frequency distributions such as symmetric and asymmetric truncated normal distributions having different widths and also for different lattice sizes. The qualitative nature of the results remains unchanged.

The second initial condition that we need to take care of is the distribution of initial phases. We will always sample the initial phases randomly from a uniform distribution but a novel feature that we will introduce in our analysis is to study the phase dynamics of the model for various widths of the initial phase distribution. As we will see, the nature of the temporal patterns changes qualitatively as we change the width of the initial phase distribution beyond a critical value. Our simulation run times will also roughly correspond to the experimental run times of 22 hrs and 36 hrs in Refs. [44, 45].

For the model parameters, we choose *K* = *a* Δ_*ω*_ as the coupling strength, where Δ_*ω*_ = 2*π* (1*/*150 −1*/*180) min^−1^ is the width of the natural frequency distribution, and vary *a* such that we are still in the weak coupling regime. The values of Λ are also chosen to satisfy this criterion while providing sufficient degree of asymmetry to the coupling function.

For all the results from numerical simulations that we will present in this paper, we have used the RK4 method [57–59] of time integration with a time step of Δ*t* = 0.01, to update the oscillator phases. The comprehensive nature of our numerical analyses is made possible by the incorporation of broadcasting and vectorization capabilities of NumPy in our code along with parallel processing capabilities on CPU servers using the multiprocessing module on Python. All of these drastically reduced computation times to the order of 100 seconds even for the large-scale simulations that generate phasemaps at different times for ranges of model parameters. This allows us to proceed with a more systematic exploration of the systems’ properties when varying multiple parameters. More details on the code are provided in the Supplement [52]. The code is publicly available on the Github repository:https://github.com/kaushik-roy-physics/2D-ERIC-local.

### B. A numerical *K*, Λ, phase atlas for self-organization

As shown in Fig. 13, we observe concentric phase waves at multiple foci at long times *t* = 1500 min for a range of (weak) *K* = *a*Δ_*ω*_ and Λ values. For these phase maps, we have sampled the initial phases from a uniform distribution between [ −*π/*2, *π/*2]. In the supplement, we have also provided timeshots at *t* = 1000, 1500, 2000 min, of the phasemap grids obtained by simulating the model for initial phases which are randomly sampled from uniform distributions between: [0, *π/*2], [ −*π/*4, *π*] and [ −*π, π*]. Additionally, we have provided animations for better visualization of the wave patterns and a Jupyter notebook that analyzes the dynamics in greater detail. For all these cases, we observe that the system exhibits concentric phase waves for a range of *K* and Λ values as long as the width of the initial phase distribution is less than or around the neighborhood of *π*. The onset of local synchronization is quick (see animations in Supplement [52]) with timescales similar to those observed in experiments [44]. The emergence of concentric phase waves occurs for higher values of Λ as *K* is increased. For the same ranges of *K*∼ [0.8−1.2]Δ_*ω*_ and Λ∼ [1.6− 2.0] values that display concentric wave patterns in Fig. 13, we notice a qualitative change in the phase landscape from concentric phase waves to spirals as the width of the initial phase distribution roughly exceeds a width of *π* [60].

In Figs. 2 and 3, we show these distinct phase landscapes obtained for initial phases randomly sampled from uniform distributions between [−*π/*2, *π/*2] and [−*π/*2, *π*] respectively. In both cases, the (*K*, Λ) values are chosen such that we are in the weak coupling regime and the timeshots of the phase maps are obtained at *t* = 1000, 1200, 1500, 2000 min.

**FIG. 2:**
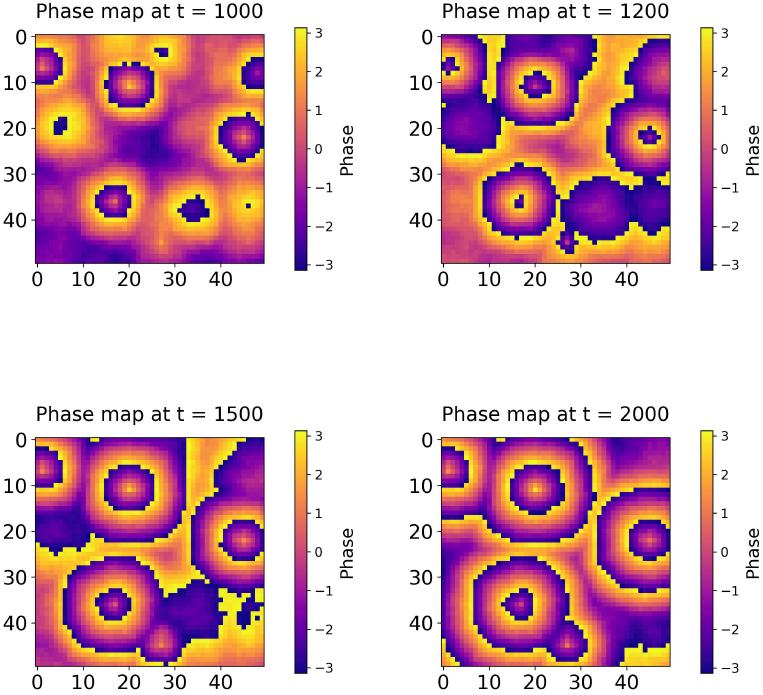
Timeshots for the phasemaps obtained by simulating Eqs. (2) to *t* = 1000, 1200, 1500, 2000 min using the RK4 method. For the initial phases, we randomly sample them from a uniform distribution between [ −*π/*2, *π/*2] and use *K* = 0.8Δ_*ω*_ min^−1^, Λ = 1.7. The final phase values are wrapped between −*π* and *π*. Even for randomly chosen initial phases from a uniform distribution of width *π*, we see concentric wave patterns emerging within the timescales of the experiments in Refs. [44, 45], and at multiple foci which are almost in phase.

**FIG. 3:**
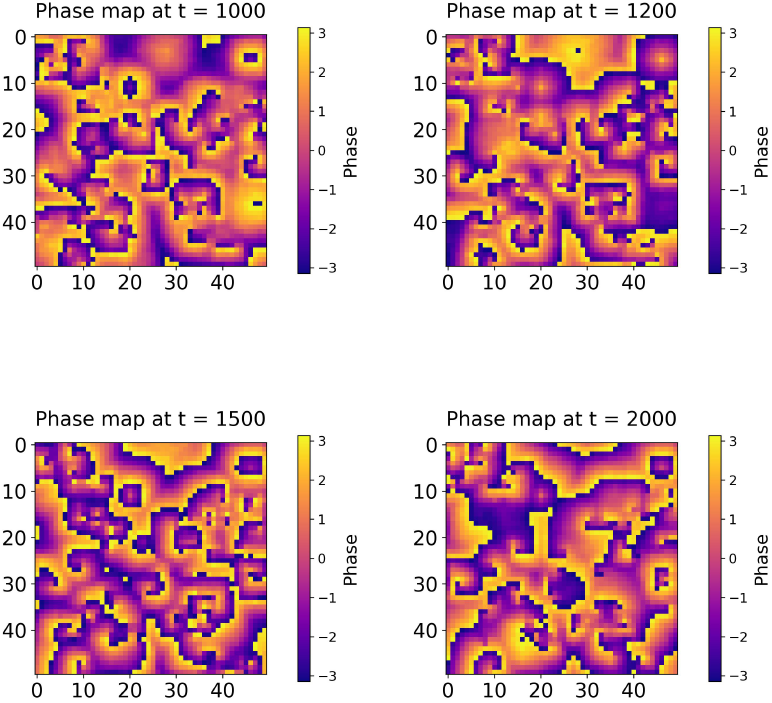
Timeshots for the phasemaps obtained by simulating Eqs. (2) to *t* = 1000, 1200, 1500, 2000 min using the RK4 method. For the initial phases, we randomly sample them from a uniform distribution between [ −*π/*2, *π*] and use *K* = 0.9Δ_*ω*_ min^−1^, Λ = 1.8. We ensure that the final phase values lie between −*π* and *π*. We see a clear distinction in the phase landscape from concentric waves to spirals as we broaden the width of the initial phase distribution.

These results are particularly encouraging for several reasons. Our model predicts concentric phase waves at multiple foci for a wide range of weak coupling strengths and ranges of randomly sampled initial phase and frequency distributions, thus consistent with the experimental observations. Importantly, we have also simulated the temporal dynamics using other canonical 2D phase models such as the original Kuramoto model, Quadratic Integrate-and-Fire (QIF) neuron models, Rectified KU-ramoto (ReKU) model (see supplement [52] for details), but none of them reproduce the wave patterns for similar ranges of natural frequencies within the timescales of our problem. Since the coupling model Eq.1 only differs from the classical Kuramoto model by the single addition of a Λ sin^2^ *θ* term, this suggests that such biharmonic coupling is necessary and sufficient to account for PSM self-organization properties.

These concentric wave patterns at multiple foci have been first observed in the randomization-dissociation experiments in Ref. [44] and also later by Hubaud et. al. [45]. Particularly noteworthy is the fact that these locally coherent spatiotemporal patterns emerge from randomized initial populations of oscillators and they are sustained under moderate variations of *K* and Λ. In biological PSM cells, *K* and Λ would correspond to the strength of the Notch-Delta coupling which depends on a variety of factors like ligand-receptor binding affinity, expression levels of Notch and Delta in the cells etc., all of which are subject to change. Yet having asymmetry and a genuine higher order harmonic in the coupling structure, such that neighboring cells affect each other’s phases differently depending on their position in the limit cycle, can explain the observation of concentric wave patterns at multiple foci under such changing conditions.

An interesting experiment that can be performed under the randomization-dissociation scenario is to broaden the width of the phase distribution at *t* = 0 of the measurements by treating the PSM cells with a Notch inhibitor (DAPT) and then allowing the phases to evolve. For sufficiently randomized initial phases, our numerical analyses predict that the dominant temporal patterns that we should observe are spirals instead of concentric waves.

### C. Emergent period

Beyond phase dynamics, the average period of coupled oscillators is strongly connected to the individual cells’ coupling properties. For instance, it is well-known that delays in weak couplings lead to an overall faster period in Kuramoto-based models [32]. Thus it is important to study how the period of the emergent process is influenced by the coupling.

Remarkably, we observe that the generation of local pacemakers under the influence of the biharmonic coupling function leads to a tightening of the average period relative to the period of PSM cells in isolation. This has also been observed in experiments [44, 45] and shows the collective effect of the locally coupled PSM oscillators. In our model analysis, we chose the limits of natural frequencies of the PSM cells to roughly correspond to experimental values for isolated mouse PSM cells. Our choice of coupling strengths also ensures that we are always in the weak coupling regime so that a phase description remains valid. In Fig. 4, we present a heatmap of the longterm average period, 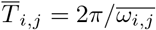 of the oscillators for the 2D ERIC model described by Eqs. (2), where the long term average frequency 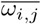 is defined as [61–64]:

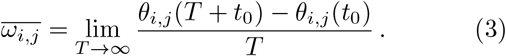

Here *t*_0_ is the transient time and *T* is the averaging time. We choose *t*_0_ = 1000 min and *T* = 1000 min meaning that we will be looking at the period averaged over roughly 6 cycles of oscillation (considering average intrinsic periods of around 150 min for the PSM cells). For the initial phases, we use the uniform distribution between [ −*π/*2, *π/*2] and the following model parameters: *K* = 0.8 Δ_*ω*_, Λ = 1.7. We observe a tightening of the long-term average period in the range [120, 165] min with the concentric foci displaying a stable period in the range [120−130] min. Thus the system self-organizes to typically follow the period of the faster oscillators in the population. This is consistent with what has been observed in the cell culture conditions recently developed [45] which allow PSM cells to exhibit stable oscillations for long periods.

**FIG. 4:**
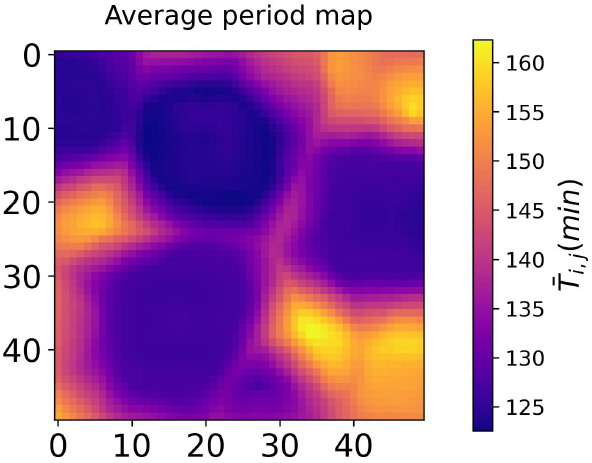
The long term average period *T* _*i*,*j*_ for the 2D ERIC model described by Eqs. (2). Note that we have sampled phases randomly from a initial phase distribution between [ −*π/*2, *π/*2]. The initial range of periods are chosen to be [150, 180] min but due to coupling effects, we see a tightening of the period in the range [120, 165] min. The foci display uniform long term average periods in the range of [120 − 130] min.

It should be noted that, unlike the experiments in [44], we do not have a time-evolving gradient in the period within the foci because we are assuming stable intrinsic oscillations for the isolated PSM cells to begin with. The establishment of a time-evolving period gradient might come from the interaction with a second oscillator, on a longer time-scale (see Refs. [28, 65]), but these interaction dynamics are not included here. However, it was noticed in [44] that reaggregates done with more anterior cells, i.e. longer oscillation periods in isolation, also self-organize with a tightening of the average period, which is consistent with our observation here that the emergent period follows the faster cells.

## III. ORIGIN OF PHASE WAVE PATTERNS IN THE 2D ERIC MODEL

A detailed mathematical understanding of the emergence of concentric phase waves at multiple foci for the biharmonic coupling function is challenging due to the finite dimensionality and the large number of oscillators, which prevents the application of methods from classical statistical mechanics. Existing literature has mainly focused on chains of coupled oscillators [63, 66–70], with limited mathematical analyses for 2D arrays of locally coupled phase oscillators [71–74]. However, these studies suggest some general features that are essential for the generation of wave patterns in oscillator arrays. For example, the role of asymmetry in coupling was highlighted (see remarks in Sec. 3.1 of Ref. [74]) for the generation of target-like patterns in oscillator arrays, albeit for monoharmonic, synaptic coupling. In contrast to target waves that require frequency heterogeneities in the population, rotating waves or spirals are connected to topological phase singularities, which occur even with symmetric, diffusive coupling and for homogenous frequencies [72]. Additionally, it was shown that the presence of asymmetry in such diffusive couplings lead to twisted spirals [72, 73]. These solutions remain stable if the coupling function derivative at zero is positive, a condition met by the biharmonic coupling considered here.

We will consider a simple scenario of a single point oscillator heterogeneity in a 1D chain of *N* = 50 ERIC model oscillators and study its effects on the local phase profile. The phase dynamics evolve according to the following equations:

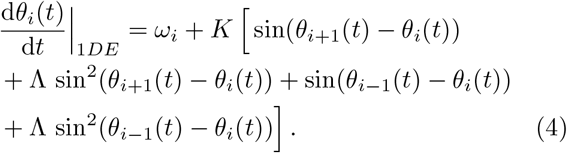

Here *θ*_*i*_(*t*) (*i* = 1, …, 50) is the phase of the *i*-th oscillator at time *t*. We will consider all oscillators to have the same (low) frequency 2*π/*180 min^−1^ except the one at *i* = 25 which has a higher frequency of 2*π/*150 min^−1^. Our choice of model parameters: *K* = 0.8Δ_*ω*_, Λ = 1.7, are also motivated by the ranges of parameters for which we observe target wave patterns for the 2D ERIC model. More importantly, the choice of coupling indicates that we have sufficient asymmetry in the coupling function and the characteristic timescales of frequency decoherence given by 1*/*Δ_*ω*_ and that of coherence (∝1*/K*) are almost similar. We will choose open boundary conditions and study the phase profile *θ*_*i*_(*t*) and neighboring phase differences *θ*_*i*+1_(*t*) −*θ*_*i*_(*t*) at *t* = 1500 min for initial phases randomly sampled from the uniform distribution between [ −*π/*2, *π/*2]. These are shown in Fig. 5. We observe a tent shaped phase profile with the sign of the phase differences changing at the location of the highest frequency oscillator.

**FIG. 5:**
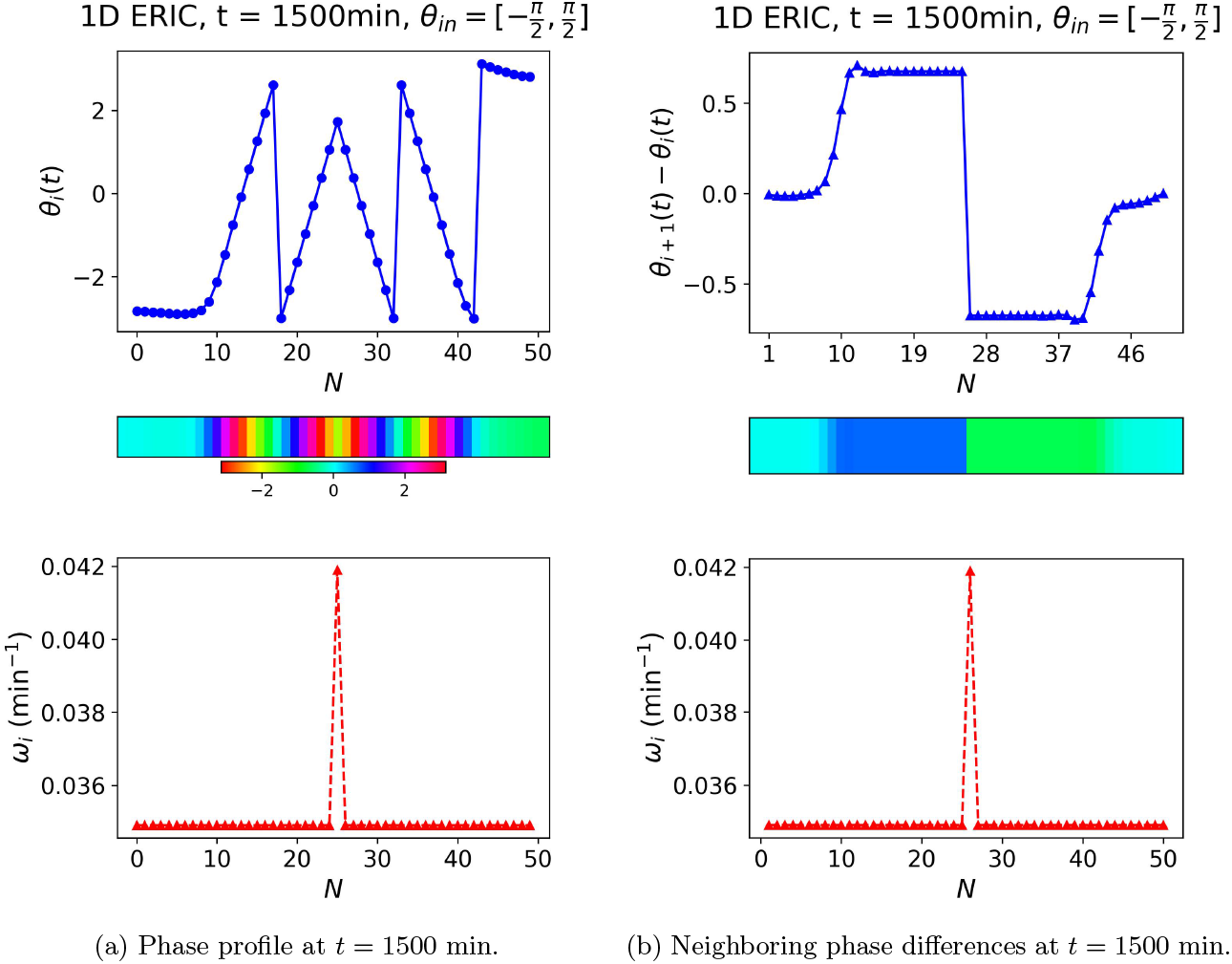
Phase profile *θ*_*i*_(*t*) and neighboring phase differences *θ*_*i*+1_(*t*) −*θ*_*i*_(*t*) (*i* = 1, …, *N* = 50) at *t* = 1500 min for the 1D chain of ERIC model oscillators interacting by Eqs. (4). The initial phases are randomly sampled from a uniform distribution between [ −*π/*2, *π/*2]. We also show the distribution of natural frequencies along the chain, in the lower graph of each figure. Notice the tent shaped phase profile around the highest frequency oscillator and the corresponding change in sign of the phase difference. The heatmap for the phase profile shown in the second row on the left figure clearly depicts the traveling wave pattern in 1D.

We will now use the same initial conditions and model parameters but consider instead a 2D array of 50×50 oscillators with the oscillator at the (*i, j*) = (25, 25) location having the highest frequency 2*π/*150 min^−1^, while others being homogenous at the frequency 2*π/*180 min^−1^. In Fig. 6, we show the 2D counterpart of the tent shaped phase profile which is a target-like wave pattern but more square shaped along the contours (see the left phasemap of Fig. 6), emanating from the highest frequency oscillator location. We have generated similar patterns at two “foci” by considering two heterogenous oscillators at two locations of the 2D array.

**FIG. 6:**
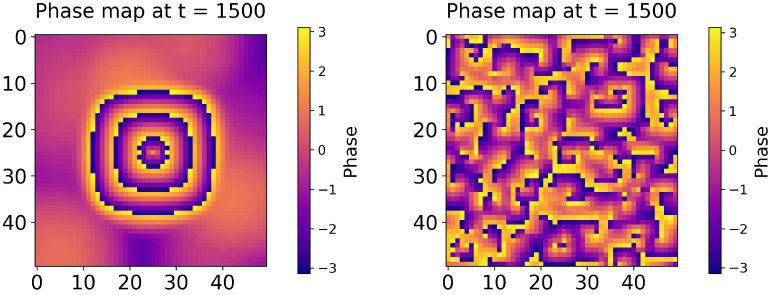
Timeshots for the phasemaps obtained by simulating the phases of the 2D ERIC model defined Eqs. (2) to *t* = 1500 min using the RK4 method. For the left figure, we randomly sample the initial phases from a uniform distribution between [ −*π/*2, *π/*2] while for the one on the right, we see a uniform distribution between [ −*π, π*]. In both cases, we use the following model parameters: *K* = 0.8Δ_*ω*_ min^−1^, Λ = 1.7. The final phase values are wrapped between −*π* and *π*. The frequencies of all the oscillators are taken constant at 2*π/*180 min^−1^, expect the one in the middle which has a frequency of 2*π/*150 min^−1^.

On the other hand, if we start with a wide distribution of initial phases chosen at random from a uniform distribution, we get twisted spirals even for a single frequency heterogeneity. The phasemap is shown on the right of Fig. 6. This demonstrates the independence of spirals on frequency heterogeneities and points towards the possibility of observing spiral waves even in homogenous PSM cell populations (such as tailbud explants) as long as we sufficiently randomize the initial phases.

A more intriguing connection regarding the role of frequency heterogeneities in generating target-like phase wave patterns emerges when considering the continuum limit [75] of the ERIC model described by Eqs. (2) (and Eqs.(4) in 1D). When phase differences are small, the continuous phase *θ*(**r**, *t*) satisfies a modified form of the non-linear phase diffusion equation given by:

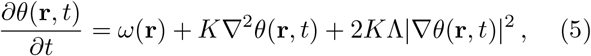

where **r** represents the coordinate vector in 2D and *ω*(**r**) is the heterogenous frequency. It should be understood that the coupling constant *K* appearing above is a rescaled version of the constant appearing in the discrete version of the phase evolution equations in Eqs. (2). Eq. (5) is known to possess target wave solutions in 2D (and traveling wave solutions in 1D) as long as the frequency heterogeneity is sufficient to destabilize the otherwise oscillatory steady state solution( see Chapter 6 of Ref.[1]). Without frequency heterogeneities and for narrow initial phase distributions, the steady-state solution is a uniformly oscillating phase, *θ*(**r**, *t*) = (*ω*_0_*/K*)*t* + *θ*(**r**, 0). For large heterogeneities *f* (**r**), defined by *ω*(**r**) = *ω*_0_ + *f* (**r**), localized regions with *f* (**r**) 0 lead to steady-state instability and target patterns emerge.

For initial phase distributions wider than *π*, the discretized 2D ERIC model cannot be expressed in the continuum form of Eq. (5). It was however shown using alternative continuum models (such as the Ginzburg-Landau equation) that the origin of spirals is independent of heterogeneities and rather dependent on initial concentrations (phase and amplitude) [1, 76]. In this model, the phase singularity occurs at the center of the spirals where the amplitude (in polar coordinates) goes to zero and as such the phase cannot be defined there.

We therefore find a generic role of symmetry breaking in local coupling and frequency inhomogeneities in 2D phase oscillator populations, in the spontaneous generation of target-like patterns. Final patterns vary with the exact form of the asymmetric coupling, model parameters, and array size. With sufficient local coupling asymmetry, higher frequency oscillators act as local pacemakers. Finally, we should caution here that there are differences between discrete and continuum cases of phase oscillator models which necessitate careful analysis, especially where a discretized description is apt. For instance, while the continuum limits of both the 2D ERIC and QIF neuron models (with coupling *h*(*θ*) = sin *θ* + Λ(1−cos *θ*)) will have the form of Eq. (5), their discrete versions exhibit distinct phase dynamics under identical parameters and initial conditions (see [52]). Understanding these behaviors requires rigorous mathematical analysis of 2D arrays, as suggested in Ref. [72, 74], to be explored in future studies.

## IV. INTRODUCING EXCITABILITY INTO THE MODEL

### A. Governing equations and phase landscapes

We will now study a simple modification of the model described by Eqs. (2) that incorporates excitability to the autonomous phase oscillators. This 2D ERIC model with excitability (2*DE* +*ex*) can be described by the following dynamical equations:

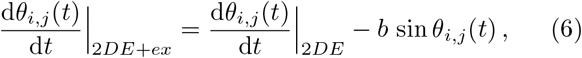

for *i, j* = 1, …, *N* and *dθ*_*i*,*j*_(*t*)*/dt* _2*DE*_ appearing on the r.h.s is given by Eqs. (2). In the absence of coupling, the isolated oscillators can be in a quiescent (for *ω*_*i*,*j*_ *< b*) or oscillatory (for *ω*_*i*,*j*_ *> b*) state. This idea is again motivated from experimental observations by Hubaud et. al. [45] which suggest that isolated PSM cells might exhibit properties of an excitable system.

In the model described above in Eqs. (6), the role of YAP is captured by the term *b*. We assume that it is roughly inversely proportional to local cell-density and saturates at high cell densities. In addition, it also depends on the cell-substrate adhesion properties at low cell-densities. For example, when we culture dissociated PSM cells at low densities in fibronectin coated substrate, both low density and cell-substrate adhesion properties lead to strong activation of YAP pathway targets resulting in a high excitability threshold (a large value of *b*). The cells are therefore in a quiescent state, *ω*_*i*,*j*_ *< b*. Replacing fibronectin with BSA leads to a decrease in *b* due to the change in cell-substrate adhesion properties which exceeds the increase due to low cell densities. In the case of explants or the randomization-dissociation experiments, the cell densities are so high that YAP signaling targets are always downregulated regardless of the substrate (*b* is always small). Hence the cells are in an oscillatory state, *ω*_*i*,*j*_ *> b* and the Notch signaling only plays a role in local synchronization of oscillations. We will now look at the phase landscapes generated under different initial conditions for phases but with the dynamics described by Eqs. (6).

As long as the cell-density is high, YAP is suppresed (*ω*_*i*,*j*_ *> b*) and we observe concentric waves at multiple foci for a wide range of *K* = *a*Δ_*ω*_ and Λ values, and for widths of the initial phase distributions close to *π*. We choose the same uniform distribution for natural frequencies as in Sec. II. In Fig. 14, we show a grid of phasemaps obtained at *t* = 1500 min by randomly sampling initial phases from a uniform distribution between [ −*π/*2, *π/*2]. We have used a broader range of *a* and Λ than the one in Fig. 13 to display the full phase landscape as a function of the coupling strengths.

As we can see, for *K* close to Δ_*ω*_ and Λ *<* 2, we have mostly target wave patterns at multiple foci; there are some instances of mixed landscapes with some spirals albeit at higher values of the coupling *K*. With an increase in asymmetry Λ in the coupling function, the landscape progressively shifts to spirals. For initial phase distributions wider than *π*, we start seeing spirals for the same range of *K* and Λ values. The contrasting pictures are shown in Figs. 7 and 8 where the timeshots of phasemaps are plotted respectively for initial phase distributions between [ −*π/*2, *π/*2] and [ −*π, π*]. The model parameters used are shown in the captions. Even though the timeshots of the phasemaps in Figs. 2 and 7 are similar, they are not identical suggesting the role of the excitable term in the dynamics.

**FIG. 7:**
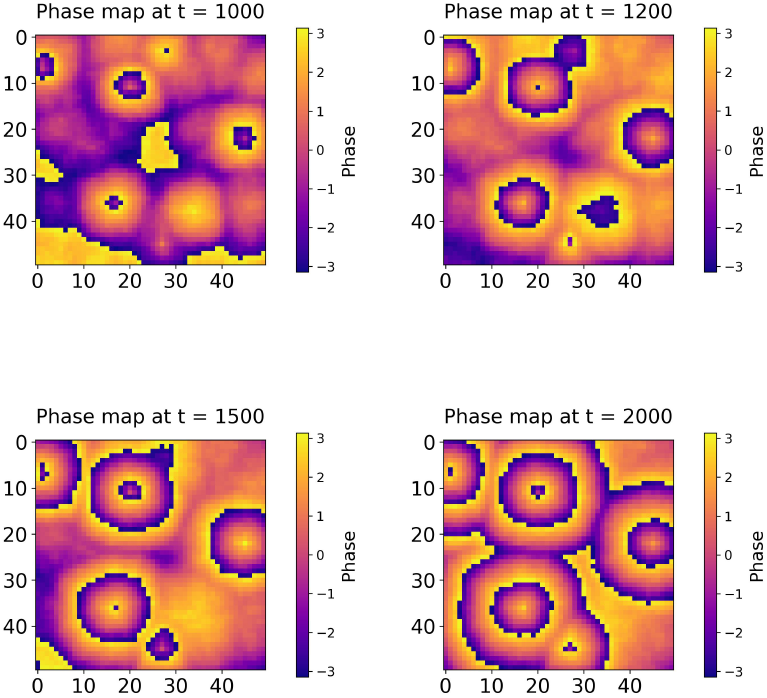
Timeshots for the phasemaps obtained by simulating Eqs. (6) to *t* = 1000, 1200, 1500, 2000 min using the RK4 method. For the initial phases, we randomly sample them from a uniform distribution between [ −*π/*2, *π/*2] and use the following model parameters: *b* = 0.01 min^−1^, *K* = 0.8Δ_*ω*_ min^−1^, Λ = 1.7. The final phase values are wrapped between −*π* and *π*. We clearly see concentric phase waves generated at three foci which are very similar (but not identical) to the *b* = 0 case.

**FIG. 8:**
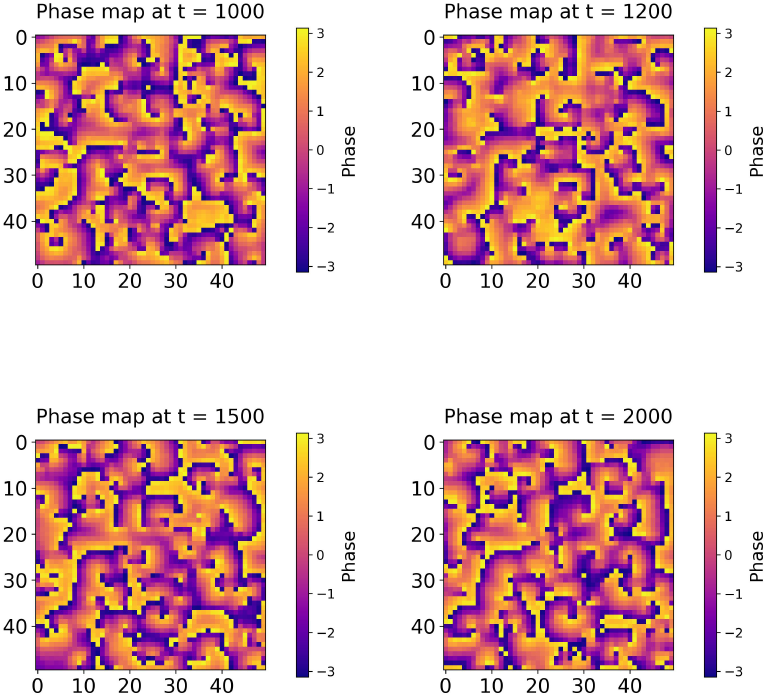
Timeshots for the phasemaps obtained by simulating Eqs. (6) to *t* = 1000, 1200, 1500, 2000 min using the RK4 method and for randomly sampled initial phases between [ −*π, π*]. We use the same model parameters: *b* = 0.01 min^−1^, *K* = 0.8Δ_*ω*_ min^−1^, Λ = 1.7 which implies that the change in phase behavior from target wave patterns to spirals is entirely due to the width of the initial phase distribution.

We can also study the behavior of the phasemaps at long times, in the (*b*, Λ) parameter space, for a given *K* close to the width of the frequency distribution. These are shown in Fig. 15 for the same choice of initial phase distribution between [ −*π/*2, *π/*2] evolved to *t* = 1500 min, using Eqs. (6), using *K* = 0.8Δ_*ω*_ and varying *b* and Λ respectively in the ranges [0−0.04] min^−1^ and [1.5− 2.0]. Note that *b* = 0 reduces the model in Eqs. (6) to the 2D ERIC model. As *b* increases (e.g for *b* = 0.036 min^−1^), we have a mixed population of quiescent and oscillatory cells for which the long term dynamics looks significantly different. There is much higher phase homogenity in the system. This can be tested experimentally by, for example, treating a fraction of cells at the beginning of the experiment with a lentiviral construct that shows high expresssion (*b* will increase) of YAP [45, 46] and using moderately high cell densities. Our analysis predicts that this would lead to disappearance of the target wave patterns, and lead to a more uniform phase structure without gradient.

### B. Local and global coherence

In this section, we will revisit the phasemaps that we obtained for the model with excitability described by Eqs. (6) and study the synchronization behavior both at the local and global level. For this, we define instantaneous local and global order parameters *ρ*_*l*:(*i*,*j*)_(*t*) and *ρ*_*g*_(*t*) respectively as:

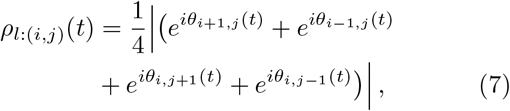

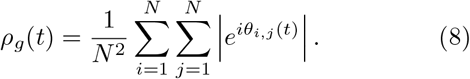

The local order parameter measures the level of phase synchronicity of the nearest neighbors at the (*i, j*)-th site. We will first discuss the local and global coherence for the case in which all the units are oscillatory to begin with i.e., *ω*_*i*,*j*_ *> b*. This is most relevant for the randomization-dissociation experiments with high cell densities that we are concerned with in this paper. We will also make predictions for the local and global coherence behavior for cases in which there is a mixed population of oscillatory and excitable elements to begin with as well as the case in which all units are excitable. As we will see, there is a marked difference in the synchronization behavior in these different cases which might be observed in experiments.

In Fig. 9, we have shown the local coherence maps at *t* = 2000 min obtained for the system in Eqs. (6) starting from initial phases randomly sampled from uniform distributions between [ −*π/*2, −*π/*2] and [ − *π, π*] respectively. For an initial phase distribution of width *π*, high local coherence develops quickly, with local partial synchrony matching the target wave patterns in Fig. 7. With a width 2*π* distribution, there is higher incoherence, shown by the blue dots on the right of Fig. 9. At these points, the phases of the neighbors are aligned almost opposite to each other in pairs resulting in a value of *ρ*_*l*_ close to zero.

**FIG. 9:**
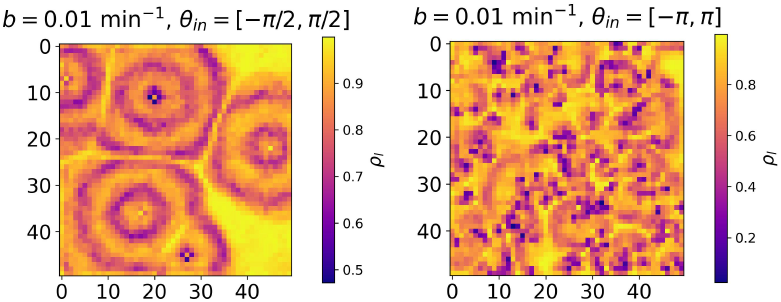
Maps of the local coherence, *ρ*_*l*:(*i*,*j*)_(*t*) at *t* = 2000 min for the system in Eqs. (6). For the initial phases, we randomly sample them from a uniform distribution between [ −*π/*2, *π/*2] (left) and [ −*π, π*] (right) respectively. We use the following model parameters: *b* = 0.01 min^−1^, *K* = 0.8Δ_*ω*_ min^−1^, Λ = 1.7.

In both cases, the system shows no sustained global coherence over time. Fig. 10 shows that starting with a narrower phase spread yields high initial global coherence, decreasing over time with damped oscillations. For a broader phase spread, global coherence increases from a disordered state but still shows oscillatory behavior. Long-term, there is no significant global phase order, due to the local nature of coupling. The oscillatory global order parameter is likely due to phase wave patterns.

**FIG. 10:**
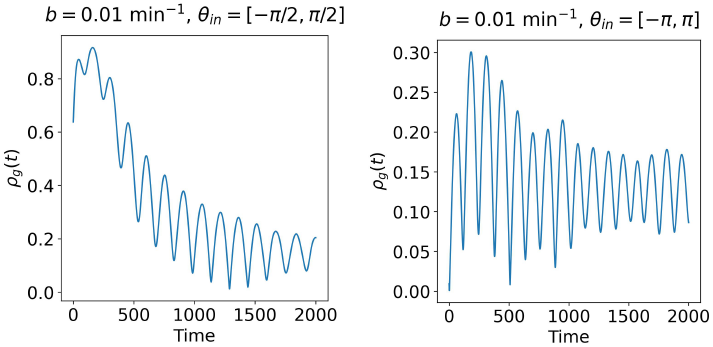
Time series of the global order parameter, *ρ*_*g*_(*t*) for initial phases randomly sampled from uniform distributions between [ −*π/*2, *π/*2] (left) and [ −*π, π*] (right) respectively. We use the following model parameters: *b* = 0.01 min^−1^, *K* = 0.8Δ_*ω*_ min^−1^, Λ = 1.7. In both cases, we see damped oscillatory behavior recapitulating the phase waves. At long times, there is no global coherence in the system for either initial condition.

Increasing the number of excitable units (higher *b*) increases global coherence over time. In Fig. 11, we show the local coherence map at *t* = 2000 min and the associated time series for the global order parameter for a mixed population of oscillatory and excitable elements (we choose *b* = 2*π/*165 min^−1^). While *ρ*_*g*_(*t*) still oscillates, there is higher global coherence. Larger *b* values result in more uniform phase structures, as seen in the phasemaps at *t* = 1500 min in Fig. 15. For intermediate *b* values, some phase waves persist, causing oscillatory global order. When *b* increases such that all *ω*_*i*,*j*_ *< b* (we choose *b* = 2*π/*145 min^−1^), the phasemaps become uniform, reflected in both the local coherence map and the global order parameter’s time series in Fig. 12.

**FIG. 11:**
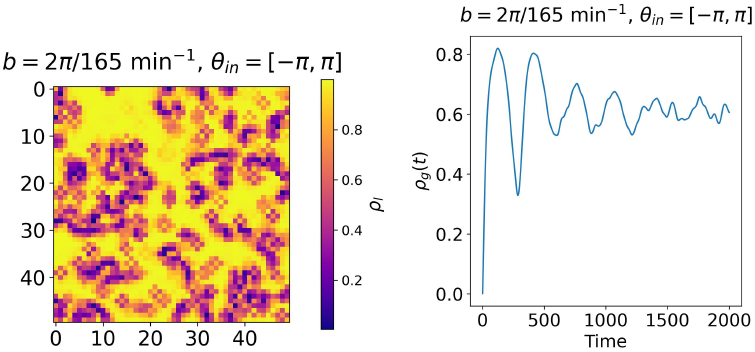
On the left, we have the local coherence map for the 2D ERIC model with excitability at *t* = 2000 min obtained for initial phases randomly sampled from a uniform distribution between [ −*π, π*]. On the right, we have the time series of the global order parameter, *ρ*_*g*_(*t*) for the same choice of initial phase distribution. In both cases, we use the following model parameters: *b* = 2*π/*165 min^−1^, *K* = 0.8Δ_*ω*_ min^−1^, Λ = 1.7. At long times, there is significant global coherence in the system.

**FIG. 12:**
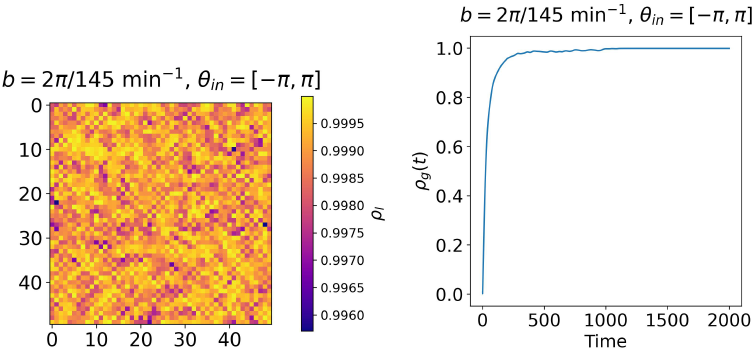
On the left, we have the local coherence map for the 2D ERIC model with excitability at *t* = 2000 min obtained for initial phases randomly sampled from a uniform distribution between [ −*π, π*]. On the right, we have the time series of the global order parameter, *ρ*_*g*_(*t*) for the same choice of initial phase distribution. In both cases, we use the following model parameters: *b* = 2*π/*145 min^−1^, *K* = 0.8Δ_*ω*_ min^−1^, Λ = 1.7. There is a high level of local synchrony in the system and a persistent global synchrony that emerges quickly after some transient time.

## V. SUMMARY AND OUTLOOK

In this work, we aim to better understand the vertebrate segmentation global oscillator by using an abstract representation of the PSM cells as phase oscillators. The essential ingredients in this analysis include an asymmetric, biharmonic coupling function, Eq. 1, and the role of initial phase and frequency distributions on the long term dynamics. This choice of the coupling differs significantly from the usual Kuramoto descriptions which consider a single sinusoidal harmonic, and is motivated by the observation of a predominantly single-sign PRC in the entrainment experiments [29], captured with the ERIC model. Additionally, we introduce excitability into the phase model and provide an interpretation of the excitability parameter in terms of the experimentally observed threshold induced by YAP signaling [45].

Our model parsimoniously explain the emergence of concentric waves with foci, in a broad range of parameters consistent with experiments. Importantly, no other simple model can explain this observation, as described in the supplement [52]. For the simple 2D Kuramoto model which includes a single sinusoidal harmonic, we do not observe any spatiotemporal pattern that resembles those observed in the mouse PSM experiments, for different initial conditions and for a wide parameter space. Similar observations are made for the Rectified KUramoto (ReKU) model which has a highly asymmetric coupling function but lacks higher order harmonics. Finally, we have considered the 2D Kuramoto model for QIF neurons which also has an asymmetric coupling function but not a genuinely higher order harmonic. This model exhibits spatiotemporal patterns but predominantly spirals and the parameter space for which we see target-like patterns is highly constrained even for initial phase distribution widths less than *π*. This is in stark contrast to our model which exhibits target patterns consistently for a wide region of the parameter space and even for initial phase distributions slightly wider than *π*. Asymmetry and the presence of higher Fourier modes are therefore essential ingredients in the coupling function for PSM cells within the phase oscillator description. One can experiment with variants of the ERIC model coupling function to find a family of coupling functions that lead to the same spatiotemporal phase dynamics.

Our simple phase models also generate concentric phase wave patterns at multiple foci for varying initial conditions of phases and natural frequencies of the phase oscillators. Notably, we hypothesize that for randomized initial populations of PSM cells distributed uniformly with widths much greater than *>> π*, we should see a departure in the phase landscape from target wave patterns into spirals. This can be experimentally tested by sufficiently decohering the phases of a population of randomized, dissociated PSM cells at the beginning of the experiment (*t* = 0) and letting them evolve. Between altering the coupling strengths, natural frequencies and initial phases, we think that tinkering with the initial phases is the most achievable and future randomization-dissociation experiments with mouse PSM cells or 2D explants can attempt to generate spirals using this method. This will further strengthen the suggestion [45] that the segmentation clock has properties of an excitable system since spirals are the signature wave patterns in such systems [8, 77–81].

Besides phase wave patterns, our phase models also explain the experimentally observed tightening of the average time period of PSM oscillations relative to their periods in isolation. This is an important validation of the asymmetric, biharmonic coupling function and highlights its role in the collective clock dynamics. We have also studied the local and global coherence patterns predicted by our excitable phase model and observed a significant difference in their behavior in the case when a fraction or all the PSM cells in the population are excitable to begin with. In experiments with explants or reaggregated dissociated cells, the high cell densities lead to lowering of the excitability threshold, *b*. Under these conditions, our model predicts concentric phase waves and these have indeed been observed in experiments [28, 44, 45]. However, if we can alter these conditions experimentally such that a fraction of the initial population stays excitable even under moderately high cell densities, our analyses suggest that we should see a decrease in phase gradients and emergence of phase homogeneity throughout the system.

Our analysis thus goes beyond the realm of modeling the experimental observations in Refs. [44, 45] but suggests novel experimental conditions that can be designed to further test the phase dynamics effected by an asymmetric, biharmonic coupling function.

Some additional features can be incorporated into our model to further capture the complexity of real biological dynamics. Firstly, we have considered a static lattice in which the cells do not move, which is fine if the timescales of cellular movement and signaling are very different from each other. If they are not, then changes in relative positions of the cells induced by cell motility can affect the signaling and hence the long time phase dynamics [82] and existing phase models of PSM cells that incorporate motility suggest that cell movement promotes global synchronization [83–87]. These models, however, use the Kuramoto form of coupling for their analysis. Ideally, we would want spatiotemporal patterns of local coherence and decoherence (see Fig. 9) to sustain even with cell movement. It would be interesting to see if this is indeed the case for our model but with mobile cells. It is also possible that there is an important role of mechanics; for example, YAP dependent mechanical cues have been found to govern the excitability threshold of the segmentation clock [45]. Additionally, in the randomization-dissociation experiments of Tsiairis et. al. [44], the ePSM reaggregates have been observed to form at characteristic distance from each other. Such regular patterns can be possible attributes of the cell-cell and cell-substrate mechanical forces [88, 89]. Lastly, in ePSM reaggregates as well as embryos, morphogen expressions change, in concordance and in response to self-organized oscillations [13]. This leads to changes of cell properties (coupling, internal frequencies) over even longer time-scales. We have not modeled such changes, keeping all parameters constant, but integrating such changes are necessary to account for long term behaviours such as wave regression and oscillation stopping [44].

To conclude, the systematic exploration of models, parameters and conditions that we provide here strongly suggests that the high order self-organization experimentally observed in randomization-dissociation experiments is a consequence of a local biharmonic, asymmetric coupling close to the one we suggest in Eq. 1. Maybe surprisingly, the study of highly perturbed biological systems, showing emergent self-organizing properties at higher scale, can thus lead to the inference of crucial properties of individual underlying components at smaller scale. This further provides an important theoretical motivation for the building and study of self-organizing embryonic systems such as the ones pioneered in [28, 44].

## ACKNOWLEDGMENTS

The work was supported by a New Frontiers in Research Fund, Exploration Grant. We thank Alexander Aulehla as well as members of the François group for useful discussions.

---

**FIG. 13:**
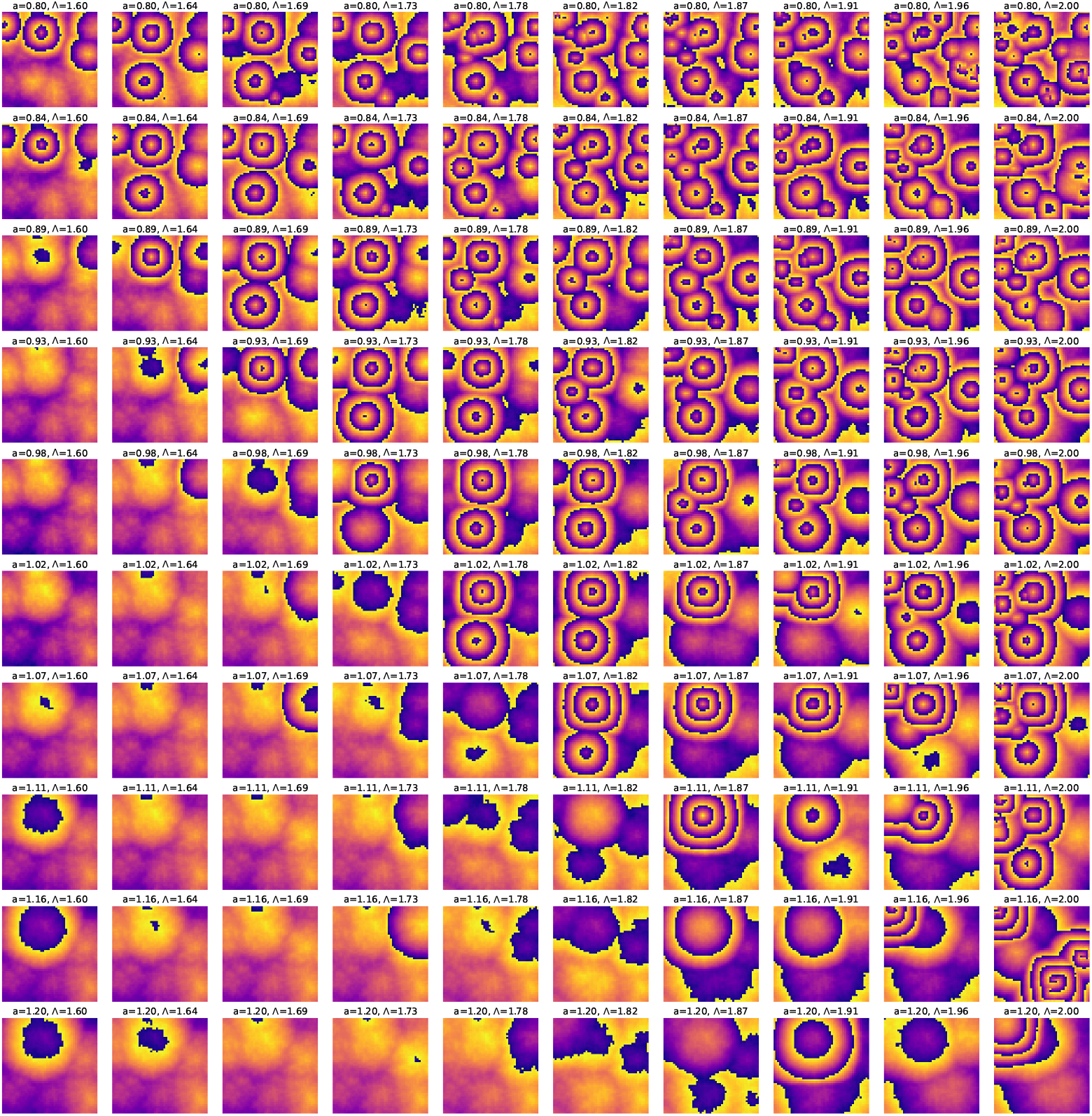
Observation of concentric phase waves for a range of coupling strengths *K* = *a*Δ_*ω*_ and Λ values. These phase maps or the heatmaps for *θ*_*i*,*j*_(*t*) (*i, j* = 1, …, 50) are obtained by simulating Eqs. (2) to *t* = 1500 min, using the RK4 time integration method with a time step of Δ*t* = 0.01. We ensure that the final phase values are wrapped between −*π* and *π*. The initial phases are randomly sampled from a uniform distribution between [ −*π/*2, *π/*2]. A noteworthy observation is that of multiple foci for phase wave generation with the number of foci roughly increasing with Λ for a given *K*.

**FIG. 14:**
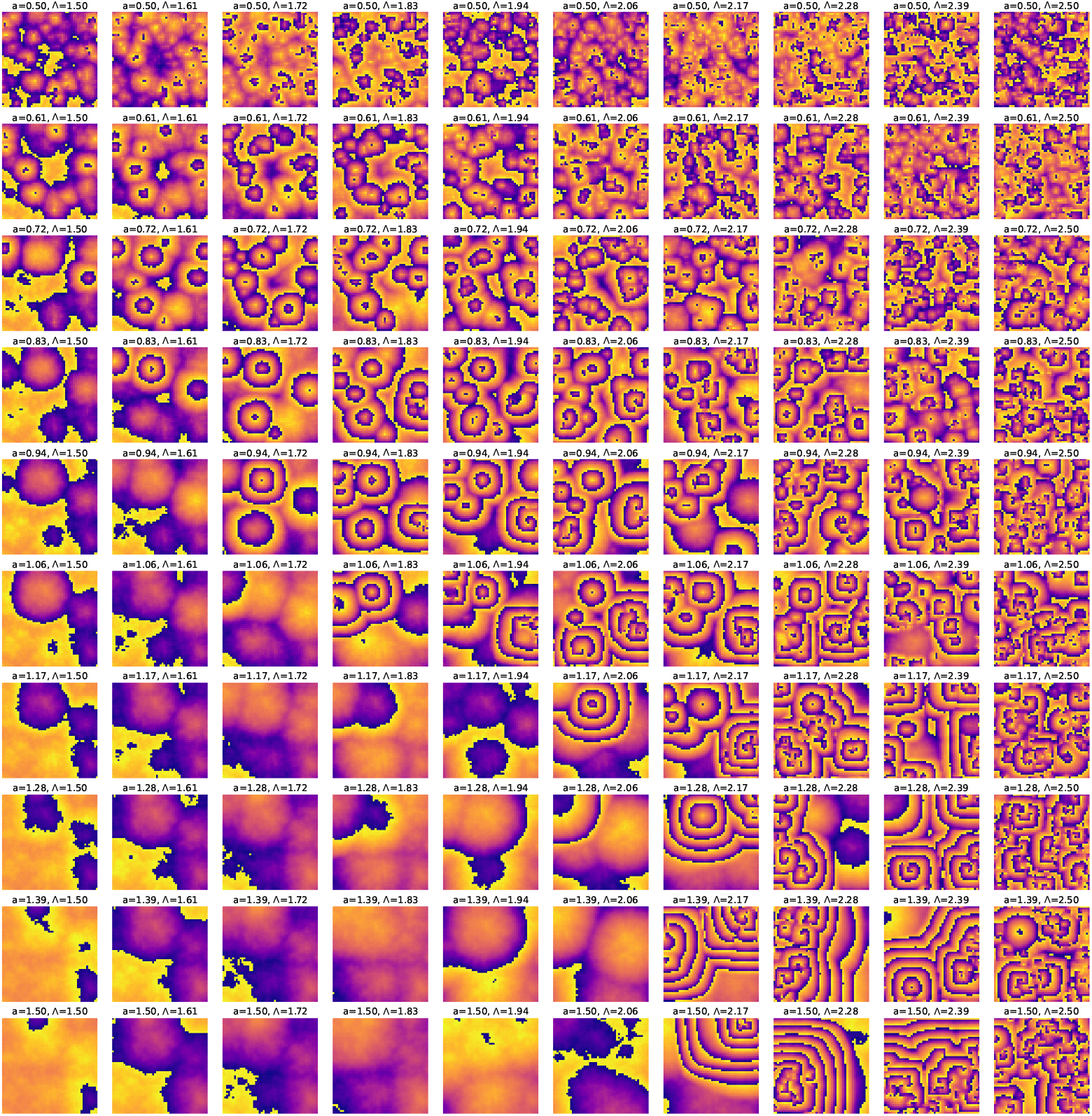
Phase landscapes at *t* = 1500 min for the locally coupled ERIC model with excitability. We use *b* = 0.01 min^−1^ for which all the uncoupled PSM cells would be in an oscillatory state. We choose 10 equally spaced values for *a* and Λ in the range [0.5 − 1.5] min^−1^ (remember *K* = *a*Δ_*ω*_) and [1.5 −2.5] respectively. The initial phases are randomly sampled from a uniform distribution between [ −*π/*2, *π/*2]. The system exhibits concentric phase wave patterns at multiple foci for a range of weak *K* and Λ values.

**FIG. 15:**
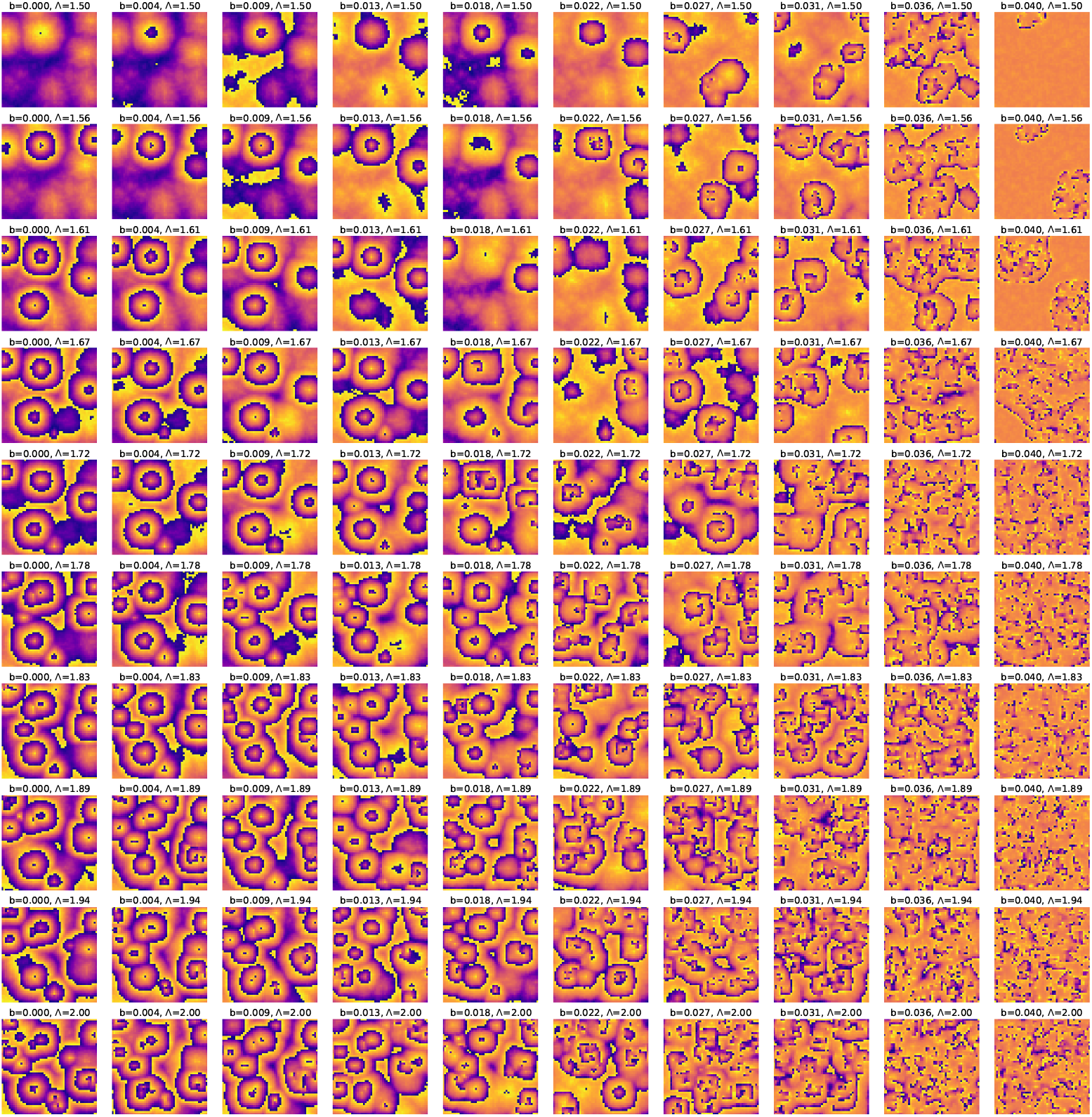
Phasemaps obtained at *t* = 1500 min for the excitable model described by Eqs. (6) for fixed *K* = 0.8Δ_*ω*_ but varying *b* and Λ in the ranges [0−0.04] min^−1^ and [1.5−2.0]. For low values of *b*, concentric wave patterns at multiple foci are observed for a wide range of Λ. As *b* increases we see a change in the temporal patterns indicating that an increase in fraction of initially excitable PSM cells can lead to disappearance of the concentric wave patterns observed in Refs. [44, 45].

## Supplemental material for

### I. DETAILS OF THE CODE

The codes for generating the plots and animations are written in Python [1] and are publicly available on the Github repository:2D-ERIC-local. We have extensively used the broadcasting and vectorization capabilities of NumPy [2] to simplify the array operations. To generate the 10×10 and 5 × 5 grids of phasemaps at different times, we have used the multiprocessing module in Python to parallelize the computation on a CPU server. The availability of the CPU server also allowed the code to run 256 processes parallely which drastically reduced the total runtime. The codes with the multiprocessing module generate multipage “.pdf” files with each page containing 100 or 25 subplots showing the phasemaps at a certain time or a fixed value of some parameter. The runtime for the entire simulation is of the order of 300 seconds for the 10 × 10 grids and 150 seconds for the 5× 5 grids. In principle, one can easily modify the code to generate phasemap grids at higher number of timepoints and for wider regions of parameter space. The computation time is not increased significantly under such a modification. We have also provided a Jupyter notebook for interactive analysis with the 2D ERIC model with or without excitability. Our code is very modular in the sense that other 2D phase models can be incorporated easily within the current implementation.

### II. PHASEMAP TIMESHOTS FOR 2D ERIC MODEL UNDER DIFFERENT INITIAL CONDITIONS

#### A. Changing initial phase distributions

In this section, we will present the timeshots, at *t* = 1000, 1500, 2000 min, of the phasemap grids generated for the 2D ERIC model given by Eq. 2 of the main text under different initial conditions. The choice of distributions for the natural frequencies and initial phases are mentioned in the figure captions. In all cases, the sampling is done at random. The phases are updated using the RK4 method of time integration with a timestep of Δ*t* = 0.01. The coupling strength is taken to be *K* = *a*Δ_*ω*_, where Δ_*ω*_ is the width of the frequency distribution. The values of *a* and Λ are also displayed in the figure labels.

In Figs. 1-3, we show the time evolution of the phasemaps from *t* = 1000 to 2000 min for initial phases which are randomly sampled from a uniform distribution between [0, *π/*2] ,i.e., of width *π/*2. The natural frequencies are randomly sampled from a uniform distribution between [2*π/*180, 2*π/*150] min^−1^. We choose a set of 100 (*a*, Λ) values by combining 10 equally spaced points in *a* ∈ [0.6− 1.2] and Λ ∈ [1.6 −2.0] respectively. We observe the formation of concentric phase waves at multiple foci for a range of (*a*, Λ) values. Similar behavior is also observed if we choose the initial phases randomly between [ −*π/*4, *π/*4] indicating that the long time phase dynamics is independent of the initial symmetry or asymmetry of the phase distribution. We also observe concentric wave patterns at multiple foci for a range of [*a*, Λ] values when we broaden the width of the initial phase distribution to *π*. One such timesnap at *t* = 1500 min is shown in Fig. 13 of the main text. Notably, we observe the possibility of concentric phase waves for initial phase distributions of width slightly greater than *π*, albeit the range of (*K* = *a*Δ_*ω*_, Λ) which allows such patterns becomes narrower. In Figs. 4-6, we have shown the time evolution of the phasemaps for the above choice of natural frequency distribution but for initial phases randomly sampled from the uniform distribution between [ −*π/*4, *π*], i.e., of width 1.25*π*.

Thus, for fairly broad uniform initial phase distributions, the 2D ERIC model with an asymmetric, biharmonic coupling function can explain the self-organization into concentric phase wave patterns at multiple foci. Only when the width of the initial phase distribution is much larger than *π*, we start seeing the dominance of spirals over target wave patterns. This theoretical prediction can be experimentally tested by randomizing the phases of the PSM cells cultured in a medium that displays stable oscillations [3] and treating them with a Notch inhibitor at the beginning of the experiment. In Figs. 8 and 9, we present the timeshots at *t* = 1000, 1500 and 2000 min of the phasemaps obtained for a wide range of (*K*, Λ) values with initial phases randomly selected from the uniform distribution of width 2*π*: [−*π, π*]. Regardless of the strength of the coupling, we always observe spirals in the phase wave patterns.

#### B. Changing natural frequency distributions

We will now fix the initial phase distribution but use different distributions for the natural frequencies and obtain the resulting phase landscapes. Instead of randomly sampling the natural frequencies from a uniform distribution, we will use symmetric and asymmetric truncated normal distributions. The rationale for this is simple: while reaggregating the dissociated cells from the PSM of several mouse embryos as in the experiments of Ref. [4], the distribution of cells might be such that the anterior of the PSM contain more cells than the posterior. This would lead to an asymmetry in the natural frequency distribution which is linked to the fraction of oscillators with frequencies within a certain range. We have analyzed several such distributions of different variances and observed concentric phase waves at multiple foci as long as the width of the initial phase distribution does not exceed *π* significantly. We have presented the phasemaps at *t* = 1000, 1500, 2000 min for two such instances of truncated normal distributions of natural frequencies in Figs. 10-12 and 13-15 respectively. In both cases, the initial phases are randomly selected from a uniform distribution between [ −*π/*2, *π/*2]. For the phasemaps in Figs. 10-12, we sample the natural frequencies randomly from a symmetric truncated normal distribution between [2*π/*180, 2*π/*150] min^−1^ with mean at 2*π/*165 min^−1^ and a variance of 2*π* (1*/*155 − 165) min^−1^. Throughout the evolution of the phases, we consistently observe concentric phase waves at multiple foci for a range of model parameters, *K* = [0.6 −1.0] Δ_*ω*_, Λ = [1.6 − 2.0]. Thus the choice of uniform distribution of frequencies is not special and the spatiotemporal phase patterns can be qualitatively reproduced by a different choice of frequency distribution. Similar results are also observed for the choice of an asymmetric truncated normal distribution between the same limits but with mean at 2*π/*170 min^−1^ and a variance of 2*π* (1*/*160−1*/*170) min^−1^. This leads to the conclusion that the consistent observation of concentric phase waves in the experiments of Refs. [3, 4] is linked to the initial phase distribution more than the distribution of natural frequencies. Since it is easier to manipulate the initial phase distribution of the PSM cells compared to their natural frequencies, we propose that experiments designed to test our predictions must tinker with the initial phase states of the PSM cells to observe a change in phase landscape from target wave patterns to spirals as predicted by our model analyses.

#### C. The case of 2D explants

Finally, we consider the case where the distribution of natural frequencies is narrow and the lattice size is also smaller. This is relevant for the case of tailbud explants which consist of only fast (high frequency) PSM cells and the total cell population is smaller compared to the case of mixing whole PSMs of several embryos. We will randomly sample frequencies from the uniform distribution between [2*π/*160, 2*π/*150] min^−1^ and initial phases from the uniform distribution between [ − *π/*2, *π/*2]. We also consider a 20×20 square lattice of phase oscillators. In Fig. 16, we have shown the grid of phasemaps at *t* = 2000 min obtained for this case. As we can see, there are target wave patterns being generated from mostly single foci for a wide range of model parameters *K* and Λ. Such patterns have been previously observed in a 2D cell culture context involving mouse tailbud explants [5] termed monolayer PSM (mPSM). Our phase model naturally explains this scenario by considering the reduction in the width of the distribution of natural frequencies (since only tailbud PSM cells are present) and decrease in the total cell population compared to the randomization-dissociation experiments.

### III. PHASEMAPS FROM ALTERNATE MODELS

In this section, we will show the phasemaps generated by other phase models which typically employ a single sinusoidal coupling. This will help distinguish the role of asymmetry in the biharmonic coupling function of the ERIC model and demonstrate the inability of other models to explain the experimentally observed temporal patterns which we are trying to model.

#### A. 2 D Kuramoto model

The phase dynamics of the 2D Kuramoto model is governed by the following equations:

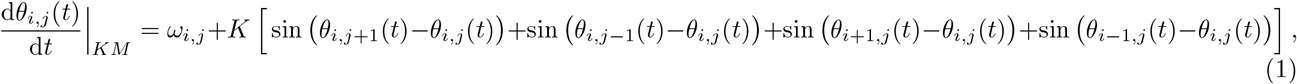

where *θ*_*i*,*j*_(*t*) represents the phase of the oscillator at (*i, j*)-th position in the 2D lattice at time *t*. We will choose open boundary conditions for the phases and evolve them using the RK4 method of time integration using time step of Δ*t* = 0.01. For the choice of natural frequency distribution, we choose the same uniform distribution between [2*π/*180, 2*π/*150] min^−1^ that we used in our main analysis. Here we show the phasemaps obtained at *t* = 1500 and 2000 min for two choices of initial phase distributions. In Fig. 17, the initial phases are randomly sampled from a uniform distribution between [ − *π/*2, *π/*2] whereas for Figs. 18 , the uniform distribution between [ −*π, π*] is used. In both cases, we use *K* = *a*Δ_*ω*_, where Δ_*ω*_ is the width of the natural frequency distribution, as the choice of coupling strength and generated phasemaps at a certain *t* for 25 equally spaced values between *a* ∈ [0.5 −3]. We can clearly observe that the 2D Kuramoto model with single sinusoidal coupling between nearest neighbors cannot generate the wave patterns within the timescales of the experiment and for a range of coupling strengths. We have even analyzed the cases with stronger coupling but observed no wave patterns for both choices of initial phase distributions.

#### B. 2 D Rectified KUramoto (ReKU) model

We will now consider a modification of the Kuramoto model with a highly asymmetric coupling function in which the neighboring oscillator phases only respond during a specific part of the cycle. In addition, the phases only speed up when they are ahead of their coupled counterparts. The model is named the Rectified Kuramoto model (ReKU) [9]; the appelation being motivated by the similarity in the functional form to the Rectified Linear Unit (ReLU) used in machine learning [10]. The coupling function is defined by:

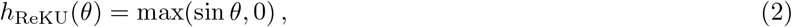

and is shown in Fig. 19 [11].

This asymmetric form of coupling is motivated by experiments performed with mixed populations of randomized PSM cells from different mouse embryos and studying their coupling behavior [9]. Real time imaging was used to quantify the phase and frequency in the mixed population and compared to independent populations. Contrary to the phase averaging prediction of the Kuramoto model, the authors observed that in the majority of cases, the cell population that is phase advanced compared to the other cell population has the ability to affect the latter’s phase. This motivated them to formulate this form of coupling which is a coarse grained version of the biharmonic coupling function in Fig. 1, in the sense that the phases do not interact at all for one half of the cycle in the ReKU model.

We will now study the phasemaps generated by such a model in 2D when the nearest neighbors are coupled to each other. The phase dynamics is governed by the following dynamical equations:

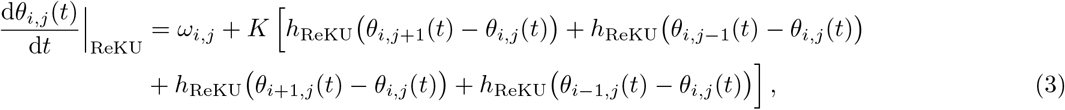

where the coupling function is defined in Eq. (2). We will sample natural frequencies randomly from the uniform distribution between [2*π/*180, 2*π/*150] min^−1^. Here we will show the phasemaps at *t* = 1000 and 2000 min for initial phases randomly sampled from uniform distributions between [ −*π/*2, *π/*2] and [ −*π, π*]. These are respectively shown in Figs. 20 and 21. We have chosen the coupling strength *K* = *a*Δ_*ω*_, where Δ_*ω*_ is the width of the frequency distribution and varied *a* between [0.5− 3.0] like in the previous section. As always, we have used open boundary conditions on the lattice and evolved the phases using the RK4 method of time integration with a time step of Δ*t* = 0.01.

With the single sinusoidal form of coupling, we observe no wave patterns for either choice of initial phases. For narrower initial phases, the phasemaps at long times show large phase homogeneities with no phase gradient, while for very wide initial phase distributions, we are likely observing topological defects as in the original Kuramoto model.

#### C. 2D Kuramoto model for populations of quadratic integrate-and-fire (QIF) neurons

We will now use a coupling of the form: sin *θ* +Λ (1 −cos *θ*), that is motivated from the Kuramoto model formulation of a population of weakly coupled QIF neurons with both electrical and chemical coupling [12]. This formulation, in turn, was motivated by a previous perturbative calculation by Izhikevich [13] involving two identical QIF neurons and also by the works in Refs. [14, 15]. In 2D, the model is described by the following dynamical equations:

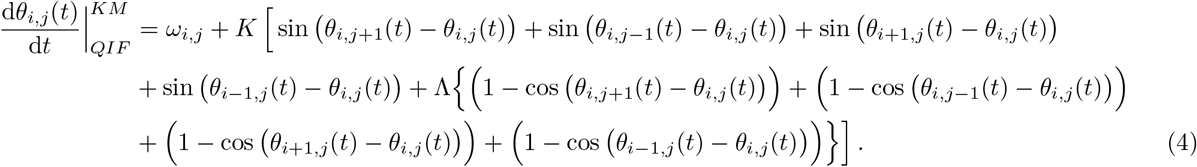

The possibility of pattern formation for the above model was studied in Ref. [16] for heterogenous lattices (non-identical frequencies) using homogenous initial conditions for the phases. The authors reported the observation of quasiregular concentric phase waves which was related to the breaking of symmetry in the coupling function due to the presence of the sin^2^(*θ/*2) term. It should be noted that the coupling in the above model is not genuinely biharmonic like the one we have used in our main analysis. We will see that this is very important to explain the experimentally observed phase patterns in addition to the requirement of asymmetry in the coupling function. In fact, it is easy to show that the coupling function of the Kuramoto model for QIF neurons can be reduced to that for the Kuramoto-Sakaguchi model [17, 18] which consists of a single sinusoidal harmonic with an overall phase shift.

We will now present the phasemaps obtained for the above model in Eqs. (4) at *t* = 1000, 1500, 2000 min for natural frequencies randomly sampled from our choice of uniform distribution between: [2*π/*180, 2*π/*150] min^−1^. We will again use different choices of initial conditions for the phases [19] For initial phases randomly sampled from a uniform distribution between [−*π/*2, *π/*2], the phasemaps are presented in Figs. 22-24 . The coupling constant is again chosen to be *K* = *a*Δ_*ω*_, where Δ_*ω*_ is the width of the frequency distribution. The values of *a* and Λ are chosen between *a* ∈ [0.5− 2.0] and Λ ∈ [1.2−1.6] respectively. We observe quasiregular concentric phase waves only for a small subset of (*K*, Λ) values. Importantly, we do not observe concentric phase waves at multiple foci indicating the importance of the higher order harmonic in the coupling function. Similar behavior is also seen for initial phase distributions of greater width such as the uniform distribution between: [ −*π/*4, *π*]. These phasemaps are shown in Figs. 25-27. Thus, asymmetry in the coupling function is important for generating phase wave patterns, specifically target wave patterns. But to consistently generate them for a range of coupling strengths and at multiple foci within the timescales of the experiment, we need to have at least a biharmonic, asymmetric coupling function. This is an important distinguishing factor between this model and the 2D ERIC model even though both have the ability to generate phase waves. Finally, for much wider initial phase distributions (figures not shown), we observe the dominance of spirals in the phasemaps at long times.

### IV. SUPPLEMENTARY MOVIES

In the folder “Animations” of the Github repository 2D-ERIC-local, we have provided the Python code for generating the movies representing phase evolution for the 2D ERIC model with and without excitability. The “.mp4” files generated by the code are also contained in the same folder. The movies are also available for viewing at the following link. For all cases, we evolve the phases to *t* = 2000 min using the RK4 method. In total, we show eight movies for the phase evolution the initial conditions and parameter choices for which are listed below:

1. 2D ERIC (2DE) model: Initial phases randomly sampled from a uniform distribution between [ −*π/*2, *π/*2], natural frequencies randomly sampled from a uniform distribution between [2*π/*180, 2*π/*150] min^−1^. The model parameters are chosen to be: *K* = 0.8 Δ_*ω*_, Λ = 1.7. Click here to download the movie.
2. 2D ERIC (2DE) model: Initial phases randomly sampled from a uniform distribution between [−*π, π*], natural frequencies randomly sampled from a uniform distribution between [2*π/*180, 2*π/*150] min^−1^. The model parameters are chosen to be: *K* = 0.8 Δ_*ω*_, Λ = 1.7. Click here to download the movie.
3. 2D ERIC model with excitability (2*DE* + *ex*): Initial phases randomly sampled from a uniform distribution between [ −*π/*2, *π/*2], natural frequencies randomly sampled from a uniform distribution between [2*π/*180, 2*π/*150] min^−1^. The model parameters are chosen to be: *K* = 0.8 Δ_*ω*_, Λ = 1.7, *b* = 0.01 min^−1^. This corresponds to a population of PSM cells in an oscillatory state prior to coupling. Click here to download the movie.
4. 2D ERIC model with excitability (2*DE* + *ex*): Initial phases randomly sampled from a uniform distribution between [−*π, π*], natural frequencies randomly sampled from a uniform distribution between [2*π/*180, 2*π/*150] min^−1^. The model parameters are chosen to be: *K* = 0.8 Δ_*ω*_, Λ = 1.7, *b* = 0.01 min^−1^. This corresponds to a population of PSM cells in an oscillatory state prior to coupling. Click here to download the movie.
5. 2D ERIC model with excitability (2*DE* + *ex*): Initial phases randomly sampled from a uniform distribution between [ −*π/*2, *π/*2], natural frequencies randomly sampled from a uniform distribution between [2*π/*180, 2*π/*150] min^−1^. The model parameters are chosen to be: *K* = 0.8 Δ_*ω*_, Λ = 1.7, *b* = 2*π/*170 min^−1^. This corresponds to a mixed population of PSM cells in excitable and oscillatory states prior to coupling. Click here to download the movie.
6. 2D ERIC model with excitability (2*DE* + *ex*): Initial phases randomly sampled from a uniform distribution between [−*π, π*], natural frequencies randomly sampled from a uniform distribution between [2*π/*180, 2*π/*150] min^−1^. The model parameters are chosen to be: *K* = 0.8 Δ_*ω*_, Λ = 1.7, *b* = 2*π/*170 min^−1^. This corresponds to a mixed population of PSM cells in excitable and oscillatory states prior to coupling. Click here to download the movie.
7. 2D ERIC model with excitability (2*DE* + *ex*): Initial phases randomly sampled from a uniform distribution between [ −*π/*2, *π/*2], natural frequencies randomly sampled from a uniform distribution between [2*π/*180, 2*π/*150] min^−1^. The model parameters are chosen to be: *K* = 0.8 Δ_*ω*_, Λ = 1.7, *b* = 2*π/*145 min^−1^. This corresponds to a population of PSM cells in excitable state prior to coupling. Click here to download the movie.
8. 2D ERIC model with excitability (2*DE* + *ex*): Initial phases randomly sampled from a uniform distribution between [−*π, π*], natural frequencies randomly sampled from a uniform distribution between [2*π/*180, 2*π/*150] min^−1^. The model parameters are chosen to be: *K* = 0.8 Δ_*ω*_, Λ = 1.7, *b* = 2*π/*145 min^−1^. This corresponds to a population of PSM cells in excitable state prior to coupling. Click here to download the movie.

---

**FIG. 1:**
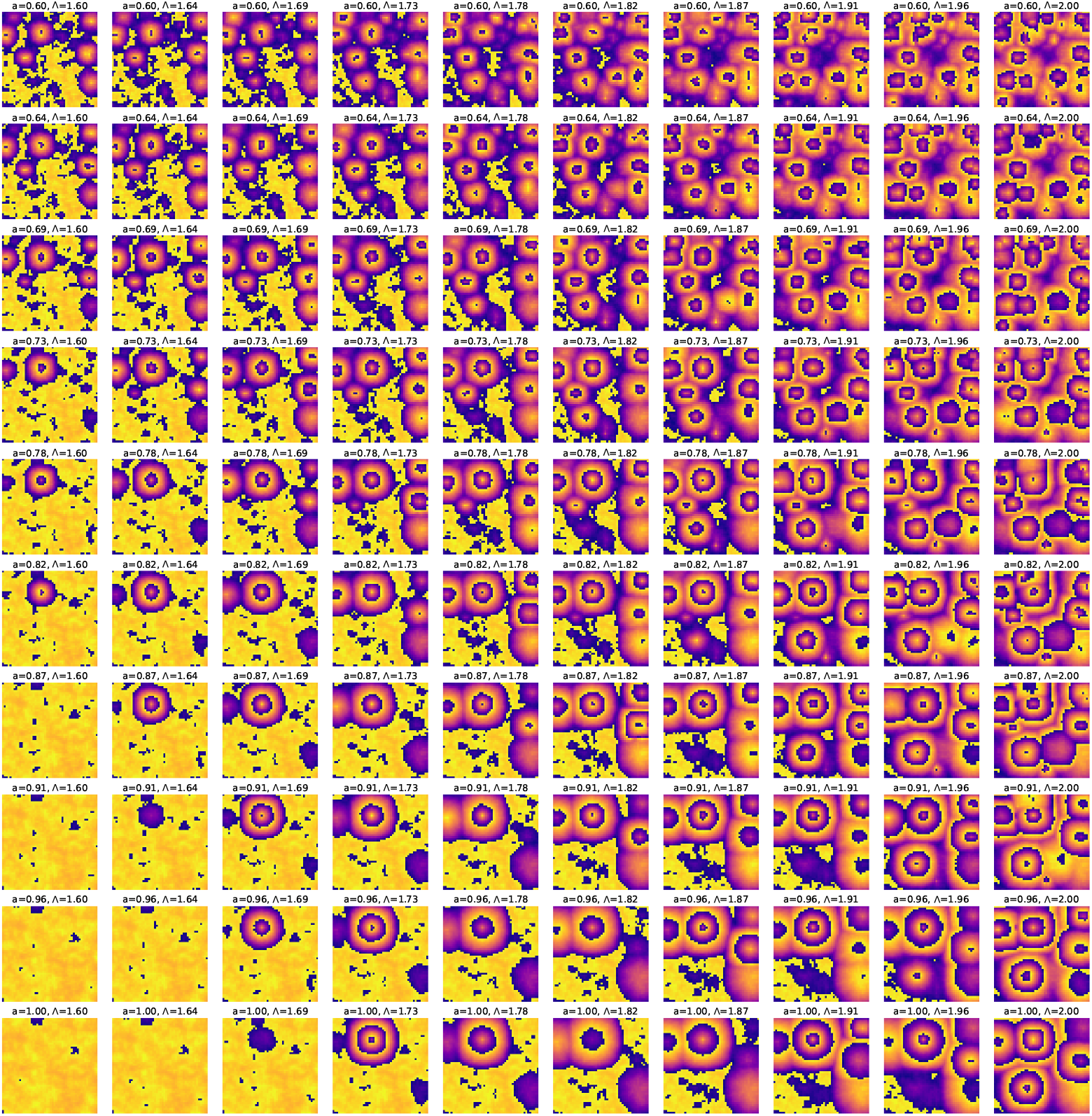
Phasemaps at *t* = 1000 min obtained by simulating the 2D ERIC (2DE) model using the RK4 method with a time step of Δ*t* = 0.01. The initial phases are randomly sampled from a uniform distribution between [0, *π/*2] and the natural frequencies are sampled randomly from a uniform distribution between [2*π/*180, 2*π/*150] min^−1^. The model parameters Λ and *K* = *a*Δ_*ω*_, where Δ_*ω*_ is the width of the frequency distribution and Λ are chosen from ten equally spaced values in Λ ∈ [1.6−2.0] and *a* ∈ [0.6−1.0]. We observe concentric phase waves at multiple foci for a lot of (*K*, Λ) values which are in the weak coupling regime of the theory.

**FIG. 2:**
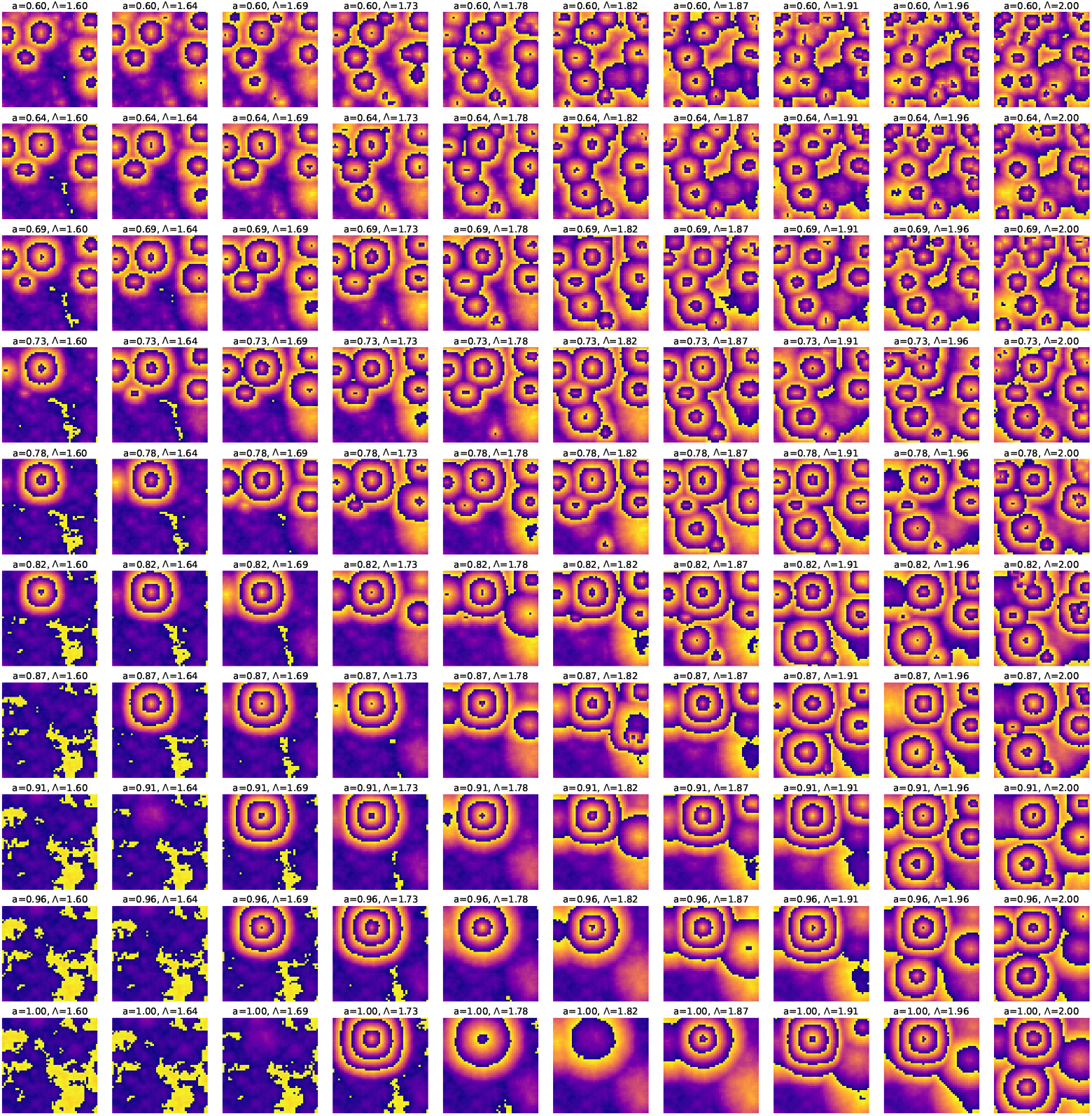
Phasemaps at *t* = 1500 min obtained by simulating the 2D ERIC (2DE) model using the RK4 method with a time step of Δ*t* = 0.01. The initial phases are randomly sampled from a uniform distribution between [0, *π/*2] and the natural frequencies are sampled randomly from a uniform distribution between [2*π/*180, 2*π/*150] min^−1^. The model parameters Λ and *K* = *a*Δ_*ω*_, where Δ_*ω*_ is the width of the frequency distribution and Λ are chosen from ten equally spaced values in Λ ∈ [1.6 −2.0] and *a* ∈ [0.6 −1.0]. We can clearly see the evolution of the phase patterns from the previous figure in Fig. 1.

**FIG. 3:**
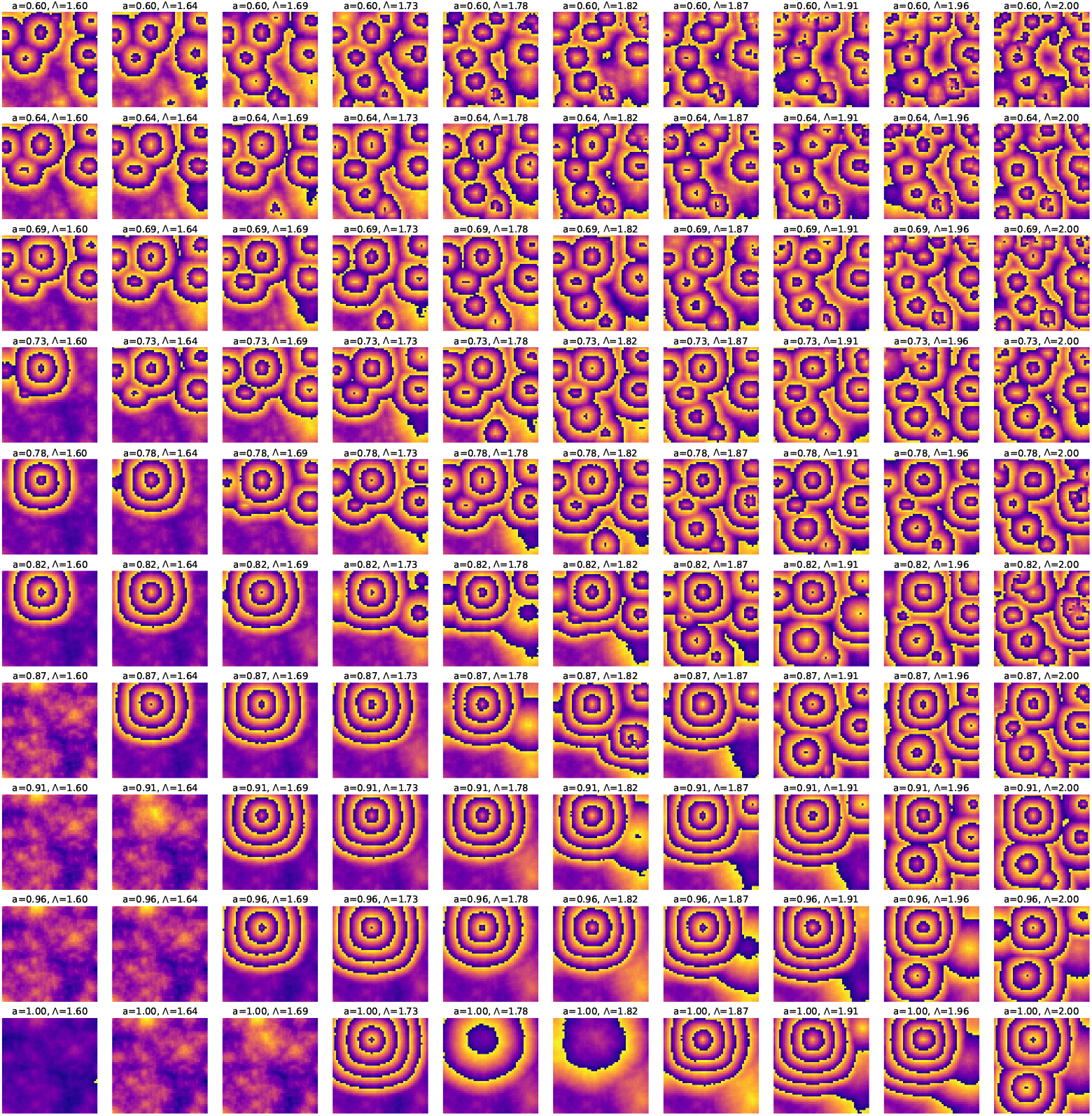
Phasemaps at *t* = 2000 min obtained by simulating the 2D ERIC (2DE) model using the RK4 method with a time step of Δ*t* = 0.01. The initial phases are randomly sampled from a uniform distribution between [0, *π/*2] and the natural frequencies are sampled randomly from a uniform distribution between [2*π/*180, 2*π/*150] min^−1^. The model parameters Λ and *K* = *a*Δ_*ω*_, where Δ_*ω*_ is the width of the frequency distribution and Λ are chosen from ten equally spaced values in Λ ∈ [1.6−2.0] and *a* ∈ [0.6 − 1.0]. The phase wave patterns sustain at long times 33 hrs which roughly correspond to the duration of the experiments in Ref. [3].

**FIG. 4:**
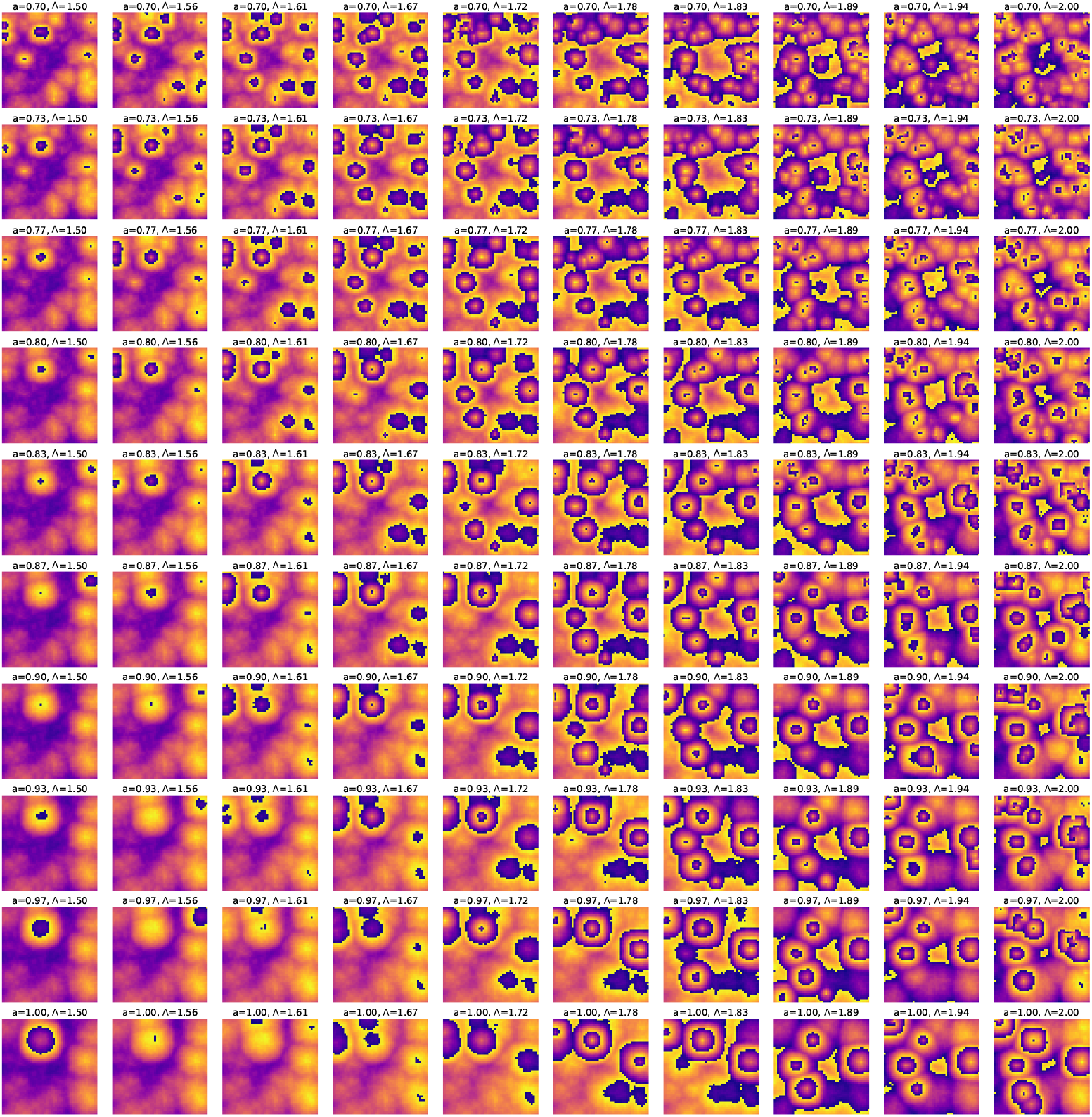
Phasemaps at *t* = 1000 min obtained by simulating the 2D ERIC (2DE) model using the RK4 method with a time step of Δ*t* = 0.01. The initial phases are randomly sampled from a uniform distribution between [−*π/*4, *π*] and the natural frequencies are sampled randomly from a uniform distribution between [2*π/*180, 2*π/*150] min^−1^. The model parameters Λ and *K* = *a*Δ_*ω*_, where Δ_*ω*_ is the width of the frequency distribution and Λ are chosen from ten equally spaced values in Λ ∈ [1.5− 2.0] and *a* ∈ [0.7 −1.0]. Even though the width of the initial phase distribution is slightly larger than *π*, we still see concentric phase waves at multiple foci for a range of (*K*, Λ). Thus, the 2D ERIC model supports the experimental observation of concentric phase waves at multiple foci for randomly sampled initial phases which are fairly widely distributed.

**FIG. 5:**
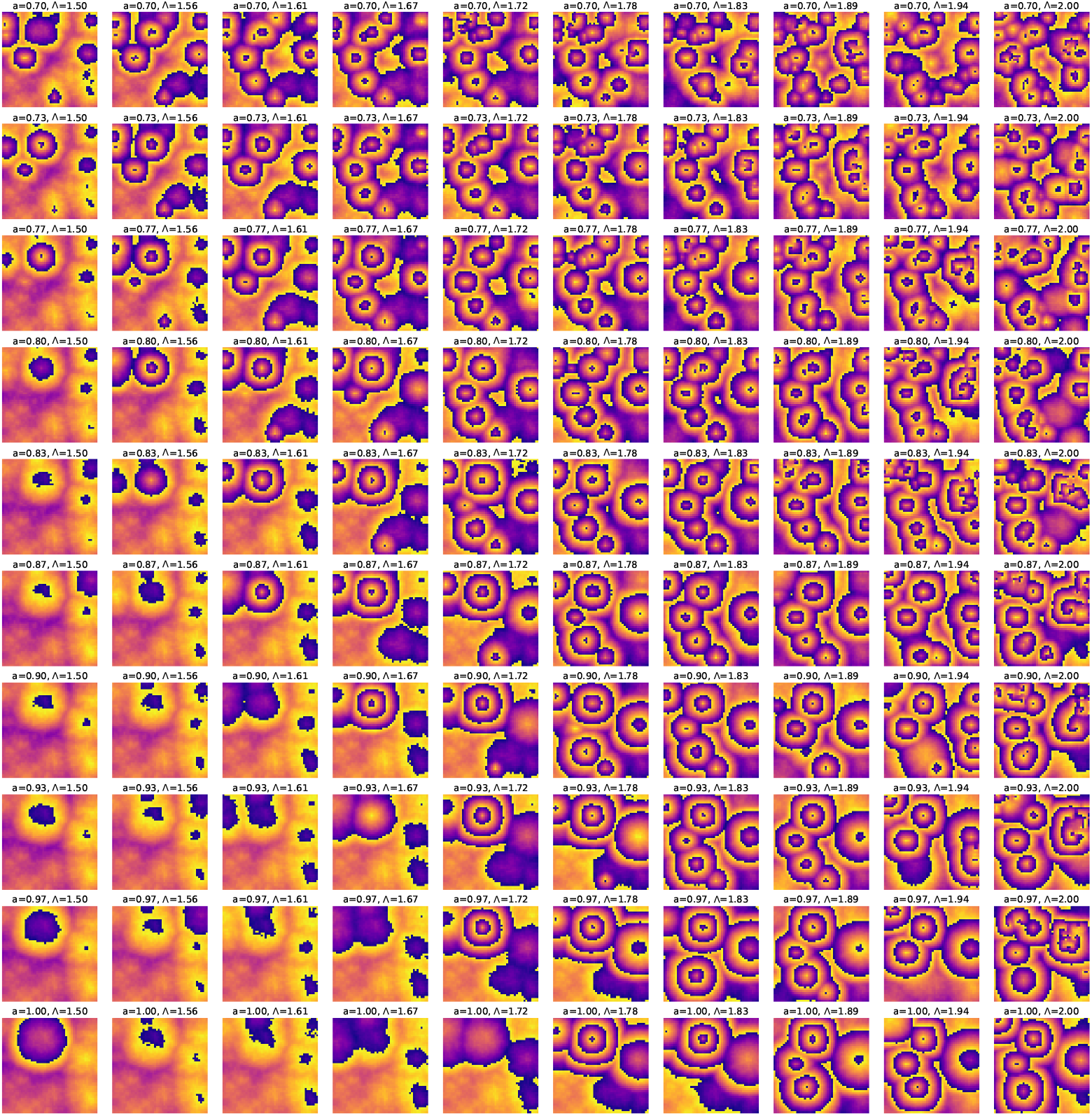
Phasemaps at *t* = 1500 min obtained by simulating the 2D ERIC (2DE) model using the RK4 method with a time step of Δ*t* = 0.01. The initial phases are randomly sampled from a uniform distribution between [−*π/*4, *π*] and the natural frequencies are sampled randomly from a uniform distribution between [2*π/*180, 2*π/*150] min^−1^. The model parameters Λ and *K* = *a*Δ_*ω*_, where Δ_*ω*_ is the width of the frequency distribution and Λ are chosen from ten equally spaced values in Λ ∈ [1.5−2.0] and *a* ∈ [0.7 −1.0]. At some large values of Λ, we obtain mixed phase landscapes that contain both concentric phase waves and spirals, but for most (*K*, Λ) values in the chosen range, we dominantly observe concentric phase waves at multiple foci.

**FIG. 6:**
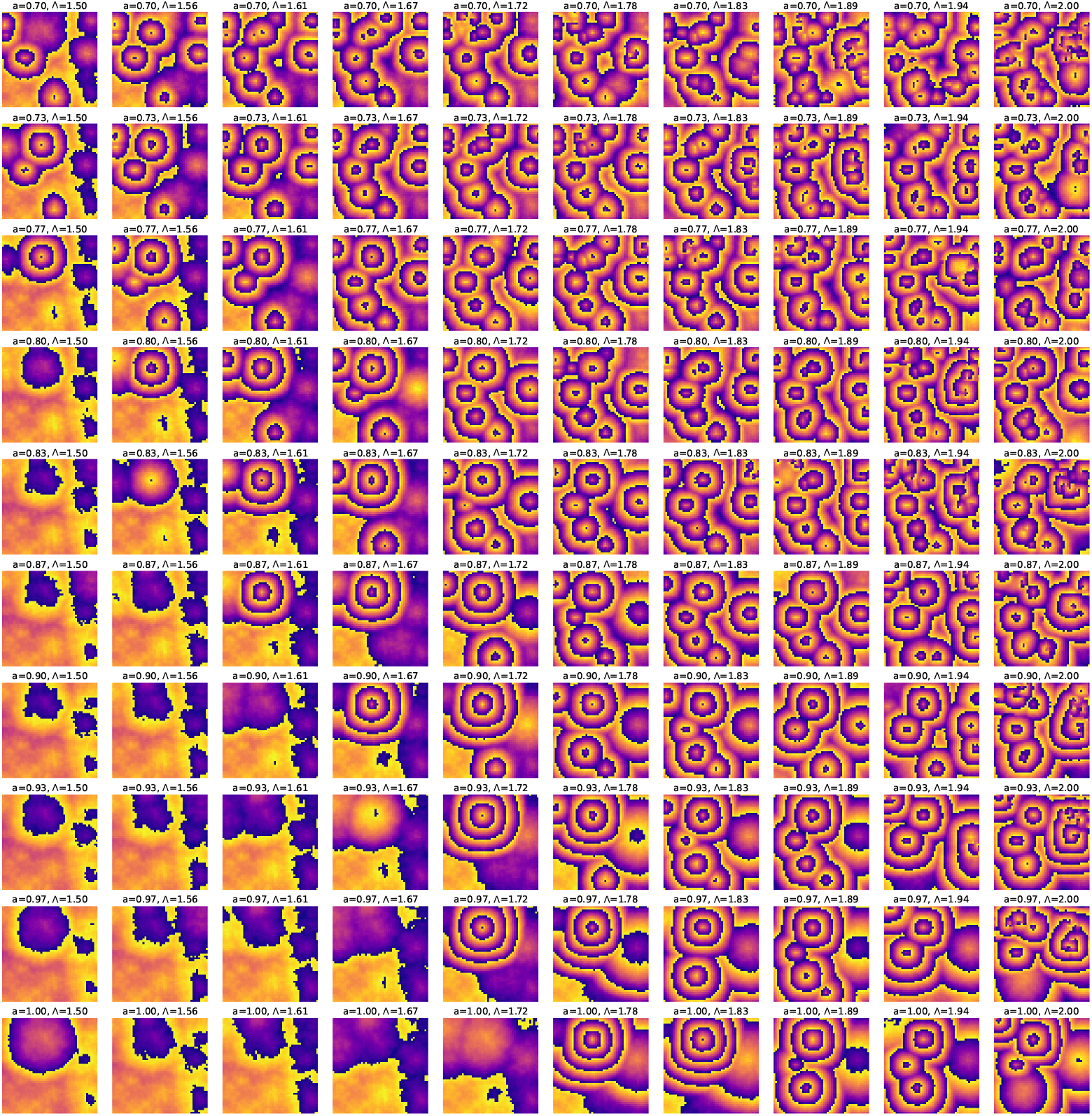
Phasemaps at *t* = 2000 min obtained by simulating the 2D ERIC (2DE) model using the RK4 method with a time step of Δ*t* = 0.01. The initial phases are randomly sampled from a uniform distribution between [ −*π/*4, *π*] and the natural frequencies are sampled randomly from a uniform distribution between [2*π/*180, 2*π/*150] min^−1^. The model parameters Λ and *K* = *a*Δ_*ω*_, where Δ_*ω*_ is the width of the frequency distribution and Λ are chosen from ten equally spaced values in Λ ∈ [1.5− 2.0] and *a* ∈ [0.7−1.0]. Like before, we see that the phase wave patterns sustain at long times.

**FIG. 7:**
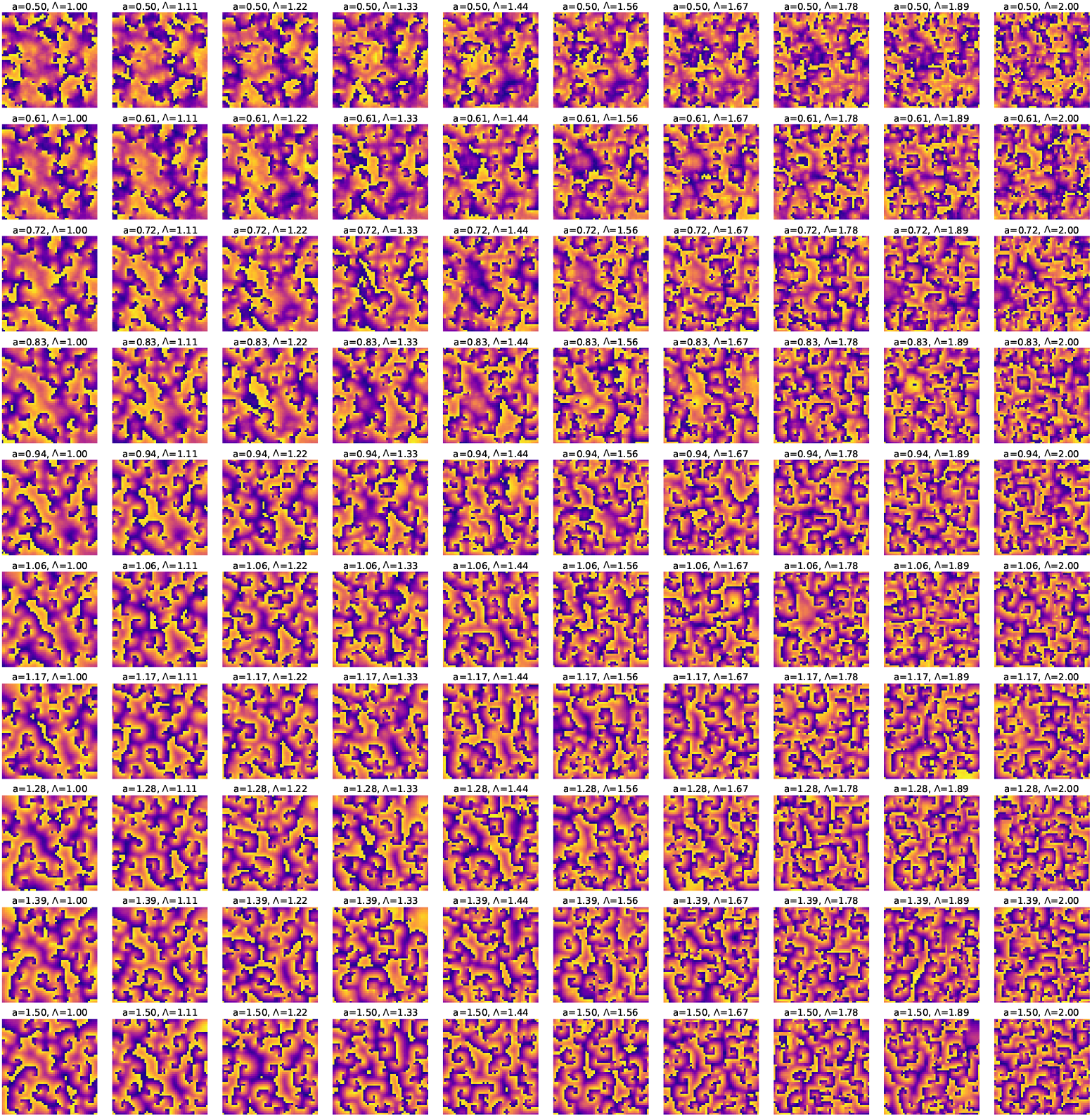
Phasemaps at *t* = 1000 min obtained by simulating the 2D ERIC (2DE) model using the RK4 method with a time step of Δ*t* = 0.01. The initial phases are randomly sampled from a uniform distribution between [ −*π, π*] and the natural frequencies are sampled randomly from a uniform distribution between [2*π/*180, 2*π/*150] min^−1^. The model parameters Λ and *K* = *a*Δ_*ω*_, where Δ_*ω*_ is the width of the frequency distribution and Λ are chosen from ten equally spaced values in Λ∈ [1.0−2.0] and *a* ∈ [0.5−1.5]. We notice a clear distinction in the phase landscape from concentric wave patterns to spirals. Note that the range of *K* and Λ is much larger compared to the previous figures.

**FIG. 8:**
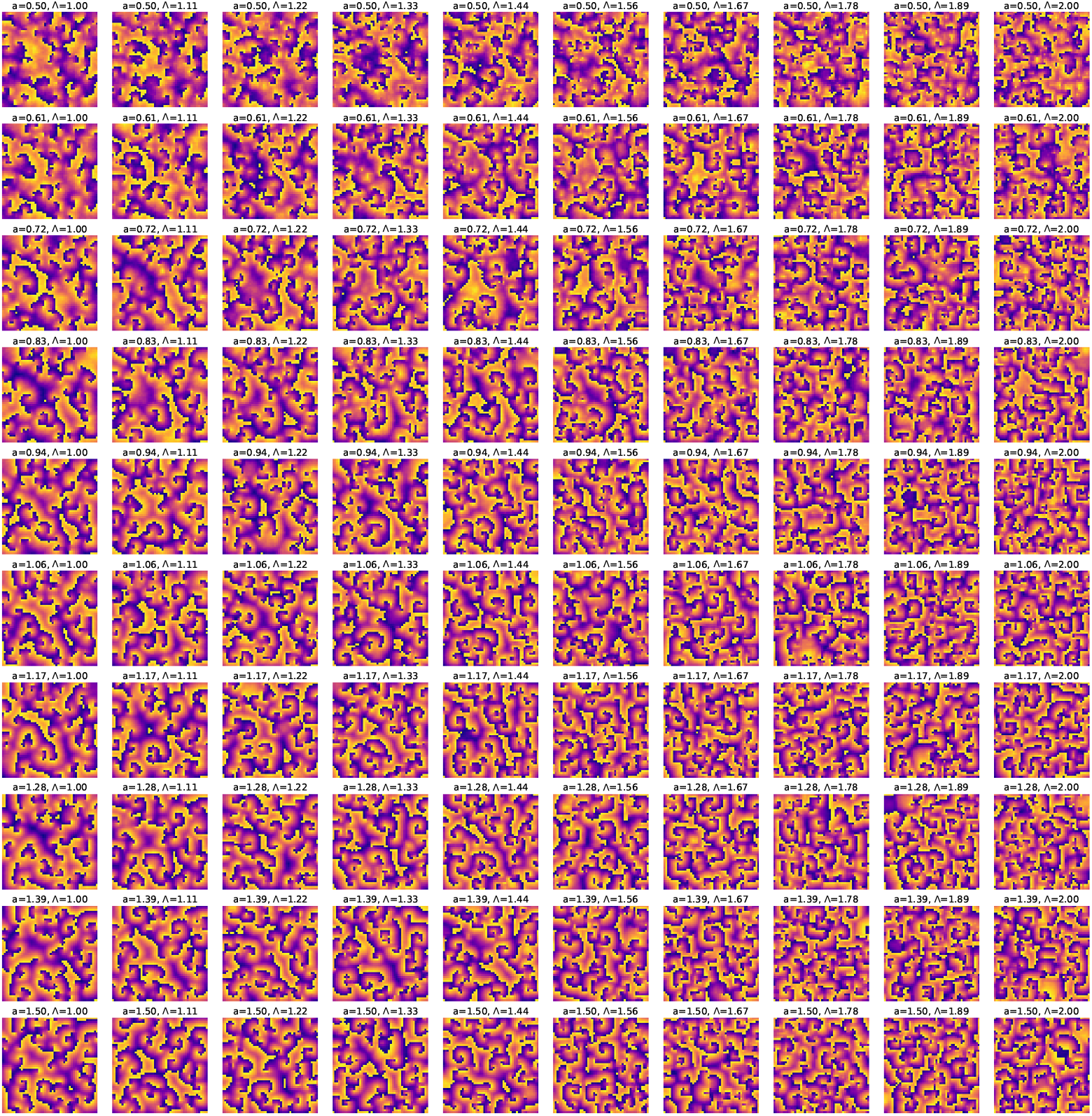
Phasemaps at *t* = 1500 min obtained by simulating the 2D ERIC (2DE) model using the RK4 method with a time step of Δ*t* = 0.01. The initial phases are randomly sampled from a uniform distribution between [−*π, π*] and the natural frequencies are sampled randomly from a uniform distribution between [2*π/*180, 2*π/*150] min^−1^. The model parameters Λ and *K* = *a*Δ_*ω*_, where Δ_*ω*_ is the width of the frequency distribution and Λ are chosen from ten equally spaced values in Λ ∈ [1.0 − 2.0] and *a* ∈ [0.5 − 1.5].

**FIG. 9:**
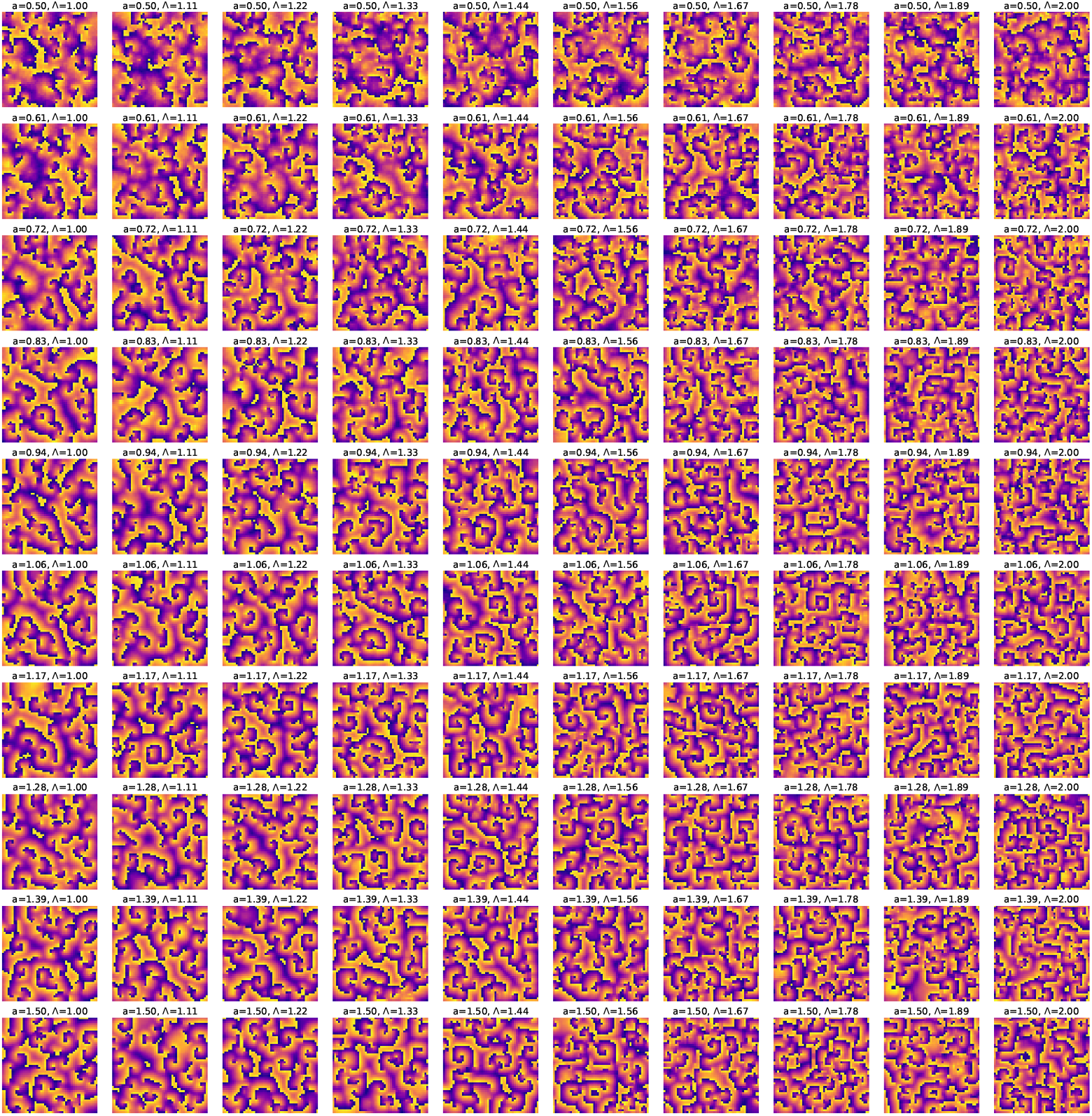
Phasemaps at *t* = 2000 min obtained by simulating the 2D ERIC (2DE) model using the RK4 method with a time step of Δ*t* = 0.01. The initial phases are randomly sampled from a uniform distribution between [−*π, π*] and the natural frequencies are sampled randomly from a uniform distribution between [2*π/*180, 2*π/*150] min^−1^. The model parameters Λ and *K* = *a*Δ_*ω*_, where Δ_*ω*_ is the width of the frequency distribution and Λ are chosen from ten equally spaced values in Λ ∈ [1.0 − 2.0] and *a* ∈ [0.5 − 1.5].

**FIG. 10:**
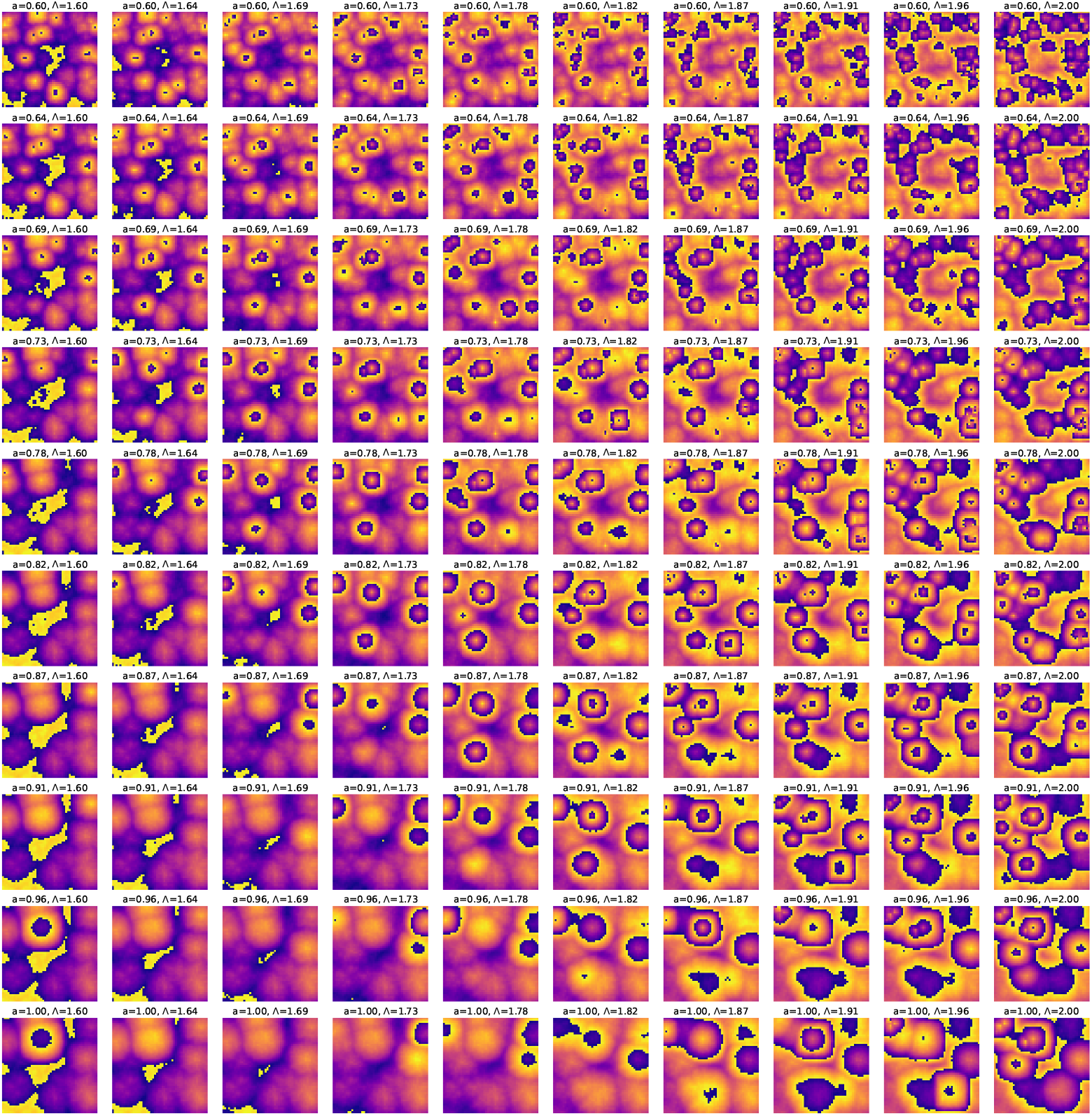
Phasemaps at *t* = 1000 min obtained by simulating the 2D ERIC (2DE) model using the RK4 method with a time step of Δ*t* = 0.01. The initial phases are randomly sampled from a uniform distribution between [ −*π/*2, *π/*2] and the natural frequencies are sampled randomly from a truncated normal distribution between [2*π/*180, 2*π/*150] min^−1^. The mean and variance of the distribution are respectively chosen to be 2*π/*165 min^−1^ and 2*π* (1*/*155 − 165) min^−1^. The model parameters Λ and *K* = *a*Δ_*ω*_, where Δ_*ω*_ is the width of the frequency distribution and Λ are chosen from ten equally spaced values in Λ ∈ [1.6 − 2.0] and *a* ∈ [0.6 −1.0]. Even for this choice of natural frequency distribution, we observe the emergence of foci for generation of concentric phase waves and for a range of (*K*, Λ) values in the weak coupling regime.

**FIG. 11:**
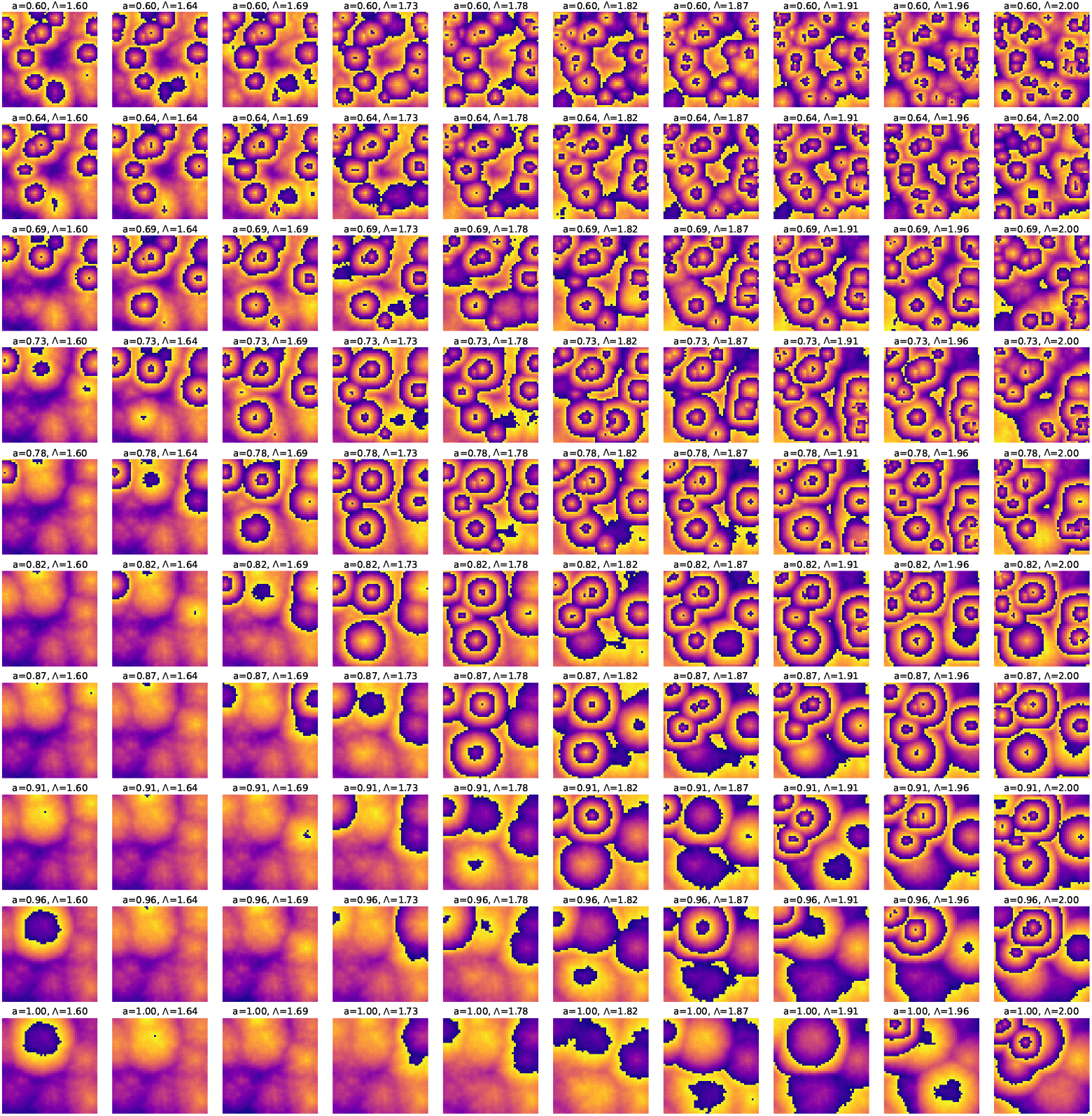
Phasemaps at *t* = 1500 min obtained by simulating the 2D ERIC (2DE) model using the RK4 method with a time step of Δ*t* = 0.01. The initial phases are randomly sampled from a uniform distribution between [ −*π/*2, *π/*2] and the natural frequencies are sampled randomly from a truncated normal distribution between [2*π/*180, 2*π/*150] min^−1^. The mean and variance of the distribution are respectively chosen to be 2*π/*165 min^−1^ and 2*π* (1*/*155 − 165) min^−1^. The model parameters Λ and *K* = *a*Δ_*ω*_, where Δ_*ω*_ is the width of the frequency distribution and Λ are chosen from ten equally spaced values in Λ ∈ [1.6 − 2.0] and *a* ∈ [0.6 − 1.0].

**FIG. 12:**
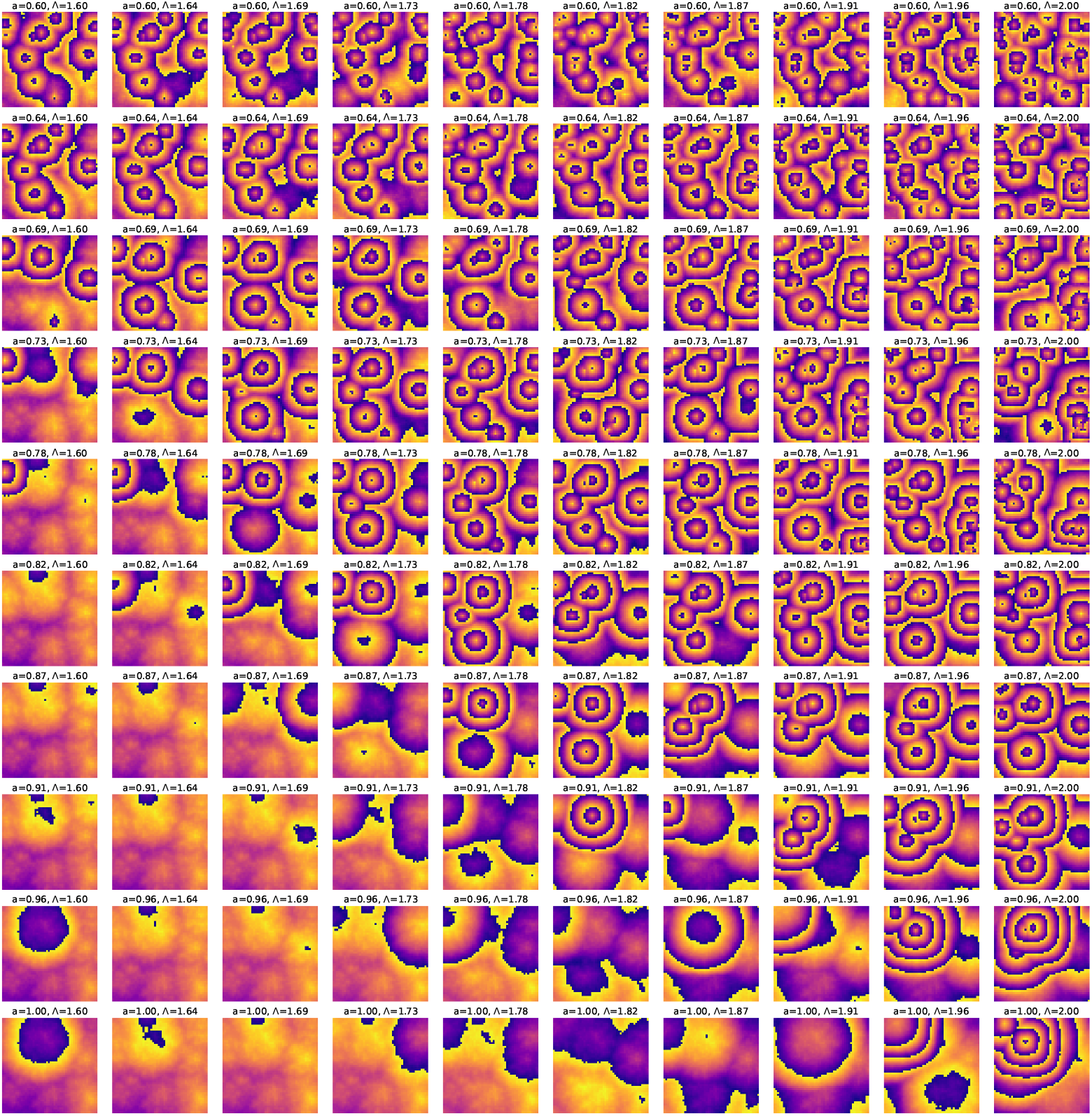
Phasemaps at *t* = 2000 min obtained by simulating the 2D ERIC (2DE) model using the RK4 method with a time step of Δ*t* = 0.01. The initial phases are randomly sampled from a uniform distribution between [ − *π/*2, *π/*2] and the natural frequencies are sampled randomly from a truncated normal distribution between [2*π/*180, 2*π/*150] min^−1^. The mean and variance of the distribution are respectively chosen to be 2*π/*165 min^−1^ and 2*π* (1*/*155 −165) min^−1^. The model parameters Λ and *K* = *a*Δ_*ω*_, where Δ_*ω*_ is the width of the frequency distribution and Λ are chosen from ten equally spaced values in Λ ∈ [1.6− 2.0] and *a* ∈ [0.6 −1.0]. For the same timescales, we see that sustained wave patterns are generated by the 2D ERIC model for natural frequencies which are not uniformly distributed but follow a symmetric truncated normal distribution.

**FIG. 13:**
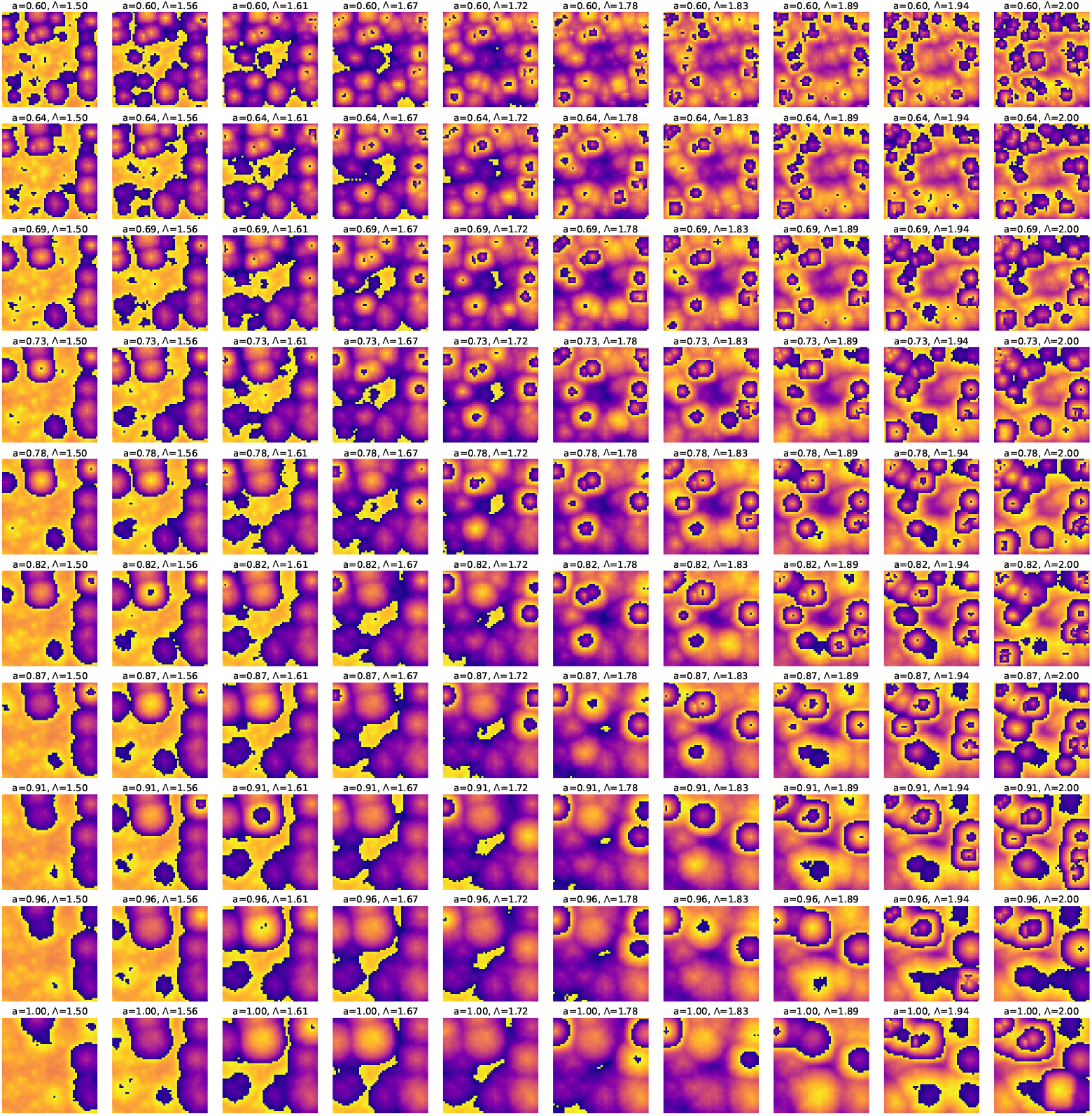
Phasemaps at *t* = 1000 min obtained by simulating the 2D ERIC (2DE) model using the RK4 method with a time step of Δ*t* = 0.01. The initial phases are randomly sampled from a uniform distribution between [ −*π/*2, *π/*2]. The natural frequencies are sampled randomly from an asymmetric, truncated normal distribution between [2*π/*180, 2*π/*150] min^−1^ with mean at 2*π/*170 min^−1^ and variance of 2*π* (1*/*160− 170) min^−1^. This corresponds to the physical case where in the randomized mixture of PSM cells, there are more cells from the anterior PSM (slowly oscillating). The model parameters Λ and *K* = *a*Δ_*ω*_, where Δ_*ω*_ is the width of the frequency distribution and Λ are chosen from ten equally spaced values in Λ ∈ [1.5 − 2.0] and *a* ∈ [0.6− 1.0]. Once again, we observe the emergence of centers for generation of concentric phase waves at weak coupling.

**FIG. 14:**
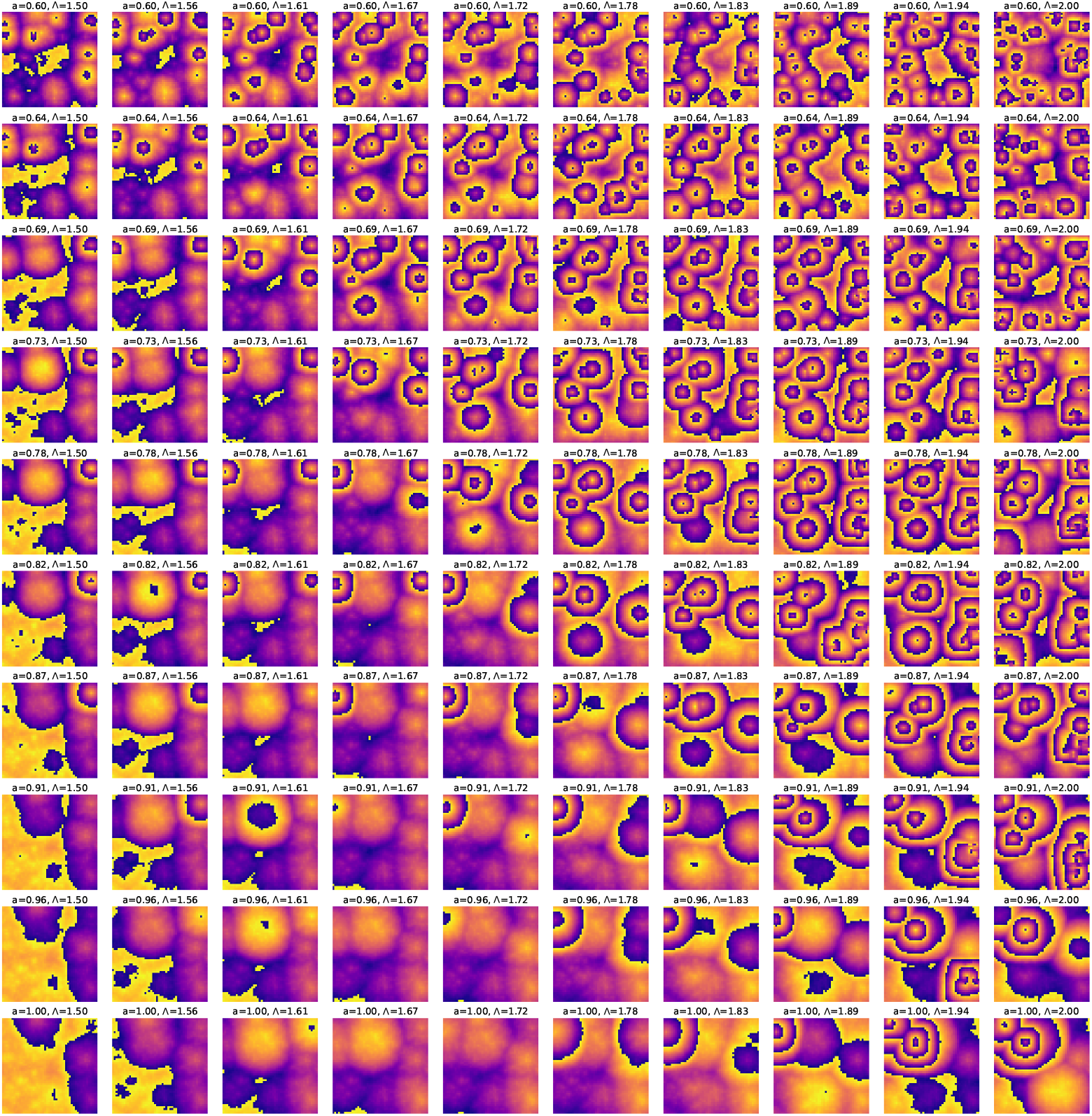
Phasemaps at *t* = 1500 min obtained by simulating the 2D ERIC (2DE) model using the RK4 method with a time step of Δ*t* = 0.01. The initial phases are randomly sampled from a uniform distribution between [ −*π/*2, *π/*2]. The natural frequencies are sampled randomly from an asymmetric, truncated normal distribution between [2*π/*180, 2*π/*150] min^−1^ with mean at 2*π/*170 min^−1^ and variance of 2*π* (1*/*160 −170) min^−1^. The model parameters Λ and *K* = *a*Δ_*ω*_, where Δ_*ω*_ is the width of the frequency distribution and Λ are chosen from ten equally spaced values in Λ ∈ [1.5 − 2.0] and *a* ∈ [0.6 − 1.0].

**FIG. 15:**
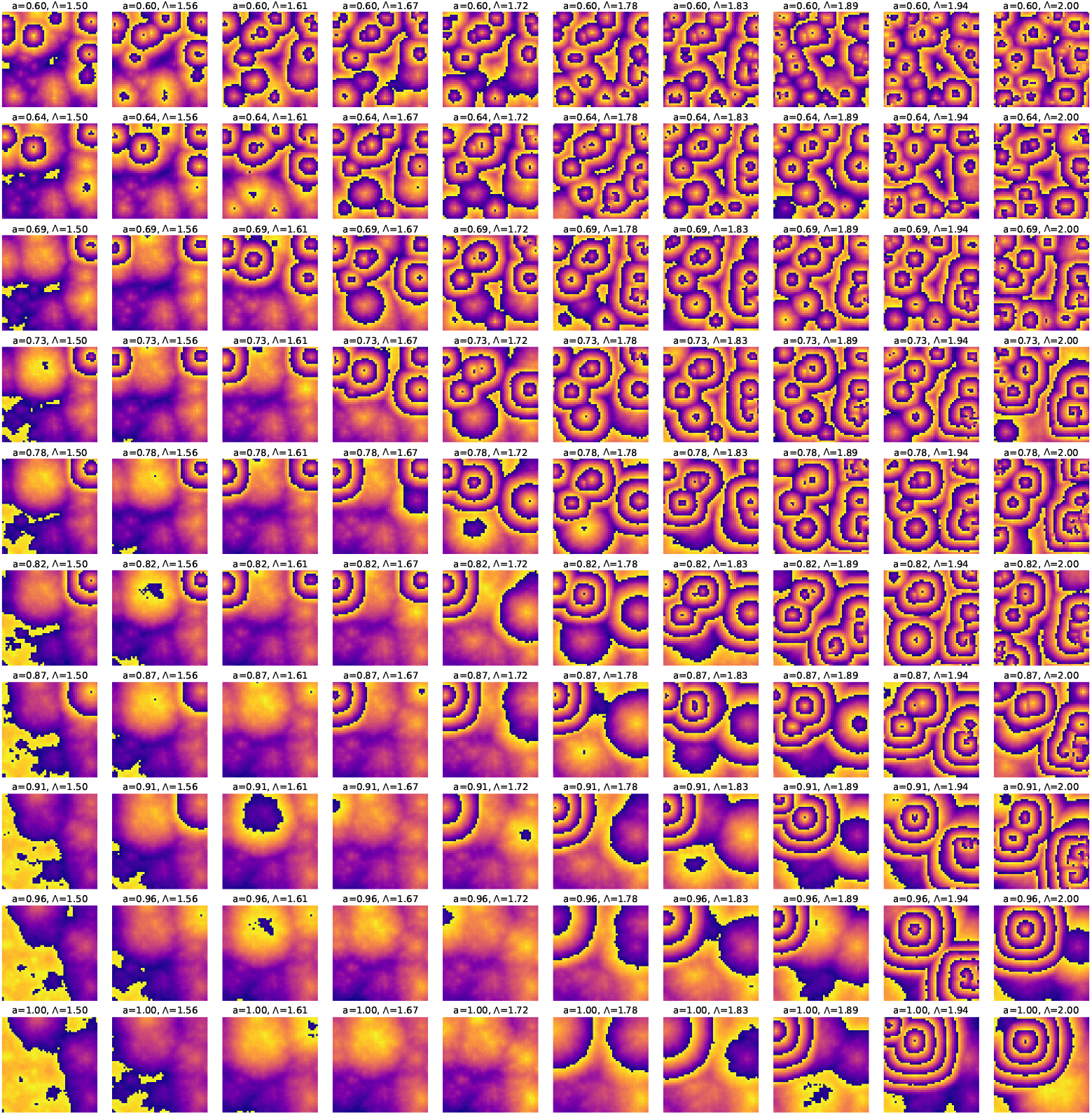
Phasemaps at *t* = 2000 min obtained by simulating the 2D ERIC (2DE) model using the RK4 method with a time step of Δ*t* = 0.01. The initial phases are randomly sampled from a uniform distribution between [−*π/*2, *π/*2]. The natural frequencies are sampled randomly from an asymmetric, truncated normal distribution between [2*π/*180, 2*π/*150] min^−1^ with mean at 2*π/*170 min^−1^ and variance of 2*π* (1*/*160 −170) min^−1^. The model parameters Λ and *K* = *a*Δ_*ω*_, where Δ_*ω*_ is the width of the frequency distribution and Λ are chosen from ten equally spaced values in Λ ∈ [1.5 − 2.0] and *a* ∈ [0.6 − 1.0]. By following the time evolution of the phasemaps, we clearly see that it is possible to generate sustained concentric phase wave patterns at multiple foci from the 2D ERIC model as long as the width of the initial phase distribution does not significantly exceed *π*.

**FIG. 16:**
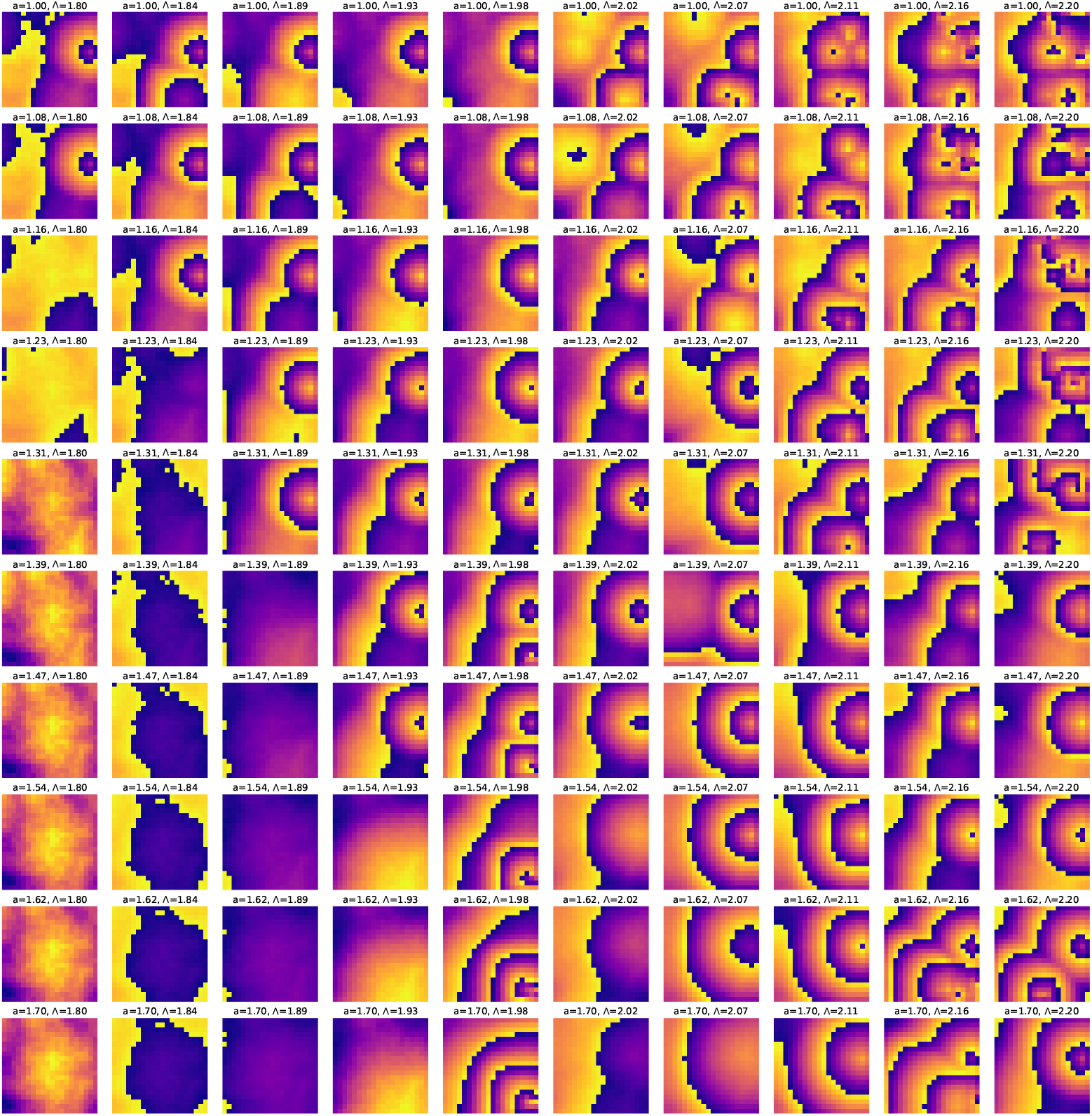
Phasemaps at *t* = 2000 min obtained by simulating the 2D ERIC (2DE) model using the RK4 method with a time step of Δ*t* = 0.01 on a 20×20 lattice. The initial phases are randomly sampled from a uniform distribution between [ −*π/*2, *π/*2]. The natural frequencies are sampled randomly from a narrow uniform distribution between [2*π/*160, 2*π/*150] min^−1^. The model parameters Λ and *K* = *a*Δ_*ω*_, where Δ_*ω*_ is the width of the frequency distribution and Λ are chosen from ten equally spaced values in Λ ∈ [1.8−2.2] and *a* ∈ [1.0− 1.7]. We observe single centers of target wave generation for the large parameter range that we have considered. This is distinct from the case of wider frequency distributions and larger lattice size where we have the possibility of target wave patterns at multiple foci.

**FIG. 17:**
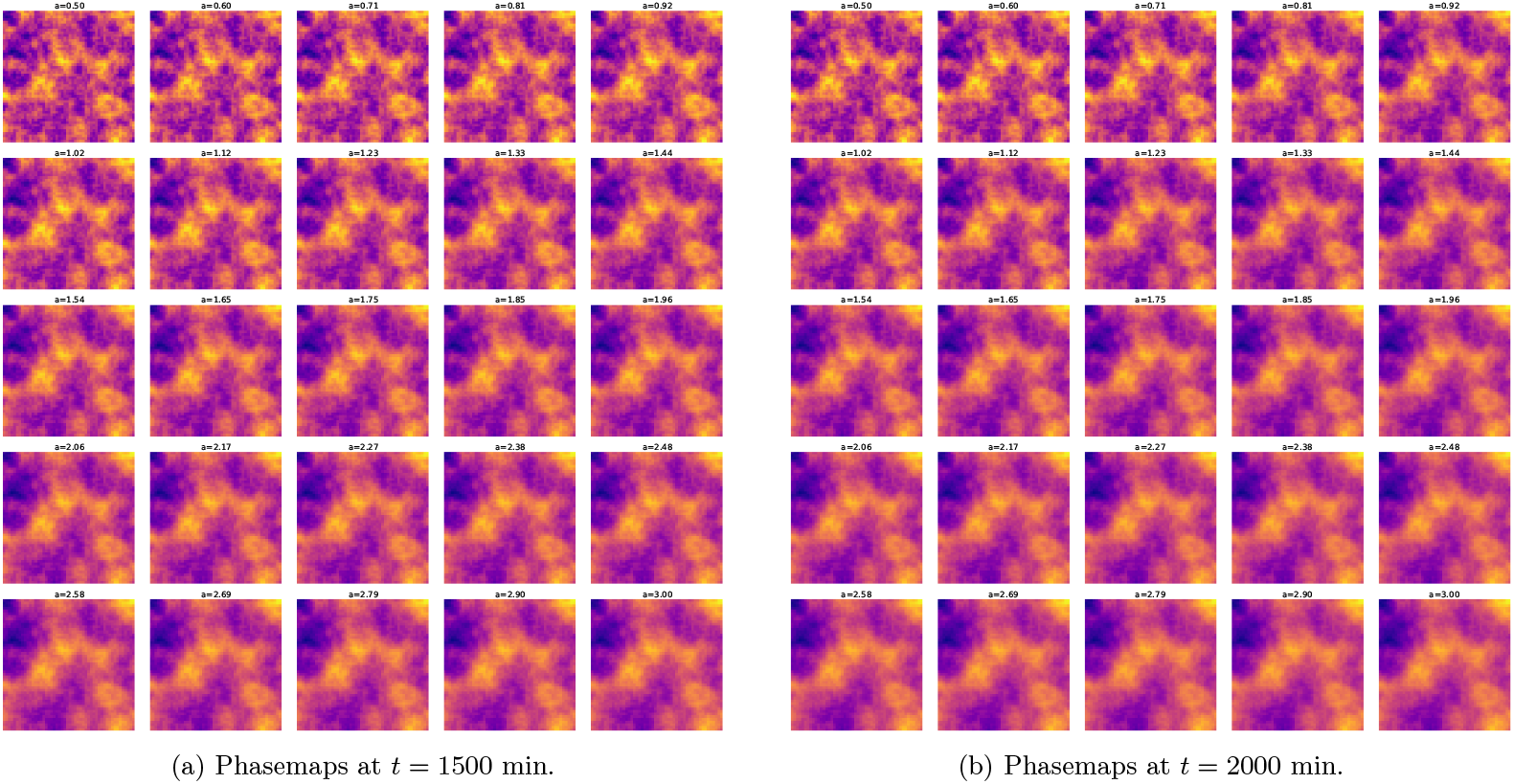
Phasemaps at *t* = 1500 and 2000 min obtained by simulating the 2D Kuramoto model with single sinusoidal coupling given by Eqs. (1). The initial phases are randomly sampled from a uniform distribution between [ −*π/*2, *π/*2]. For the natural frequencies, we randomly sample them from a uniform distribution between [2*π/*180, 2*π/*150] min^−1^, since we would like to test these canonical models against experimental frequencies of PSM cells. 25 equally spaced values for the coupling constant *K* = *a*Δ_*ω*_ are used where *a* ∈ [0.5 −3]. No wave patterns are observed within the timescales of the experiment (and beyond).

**FIG. 18:**
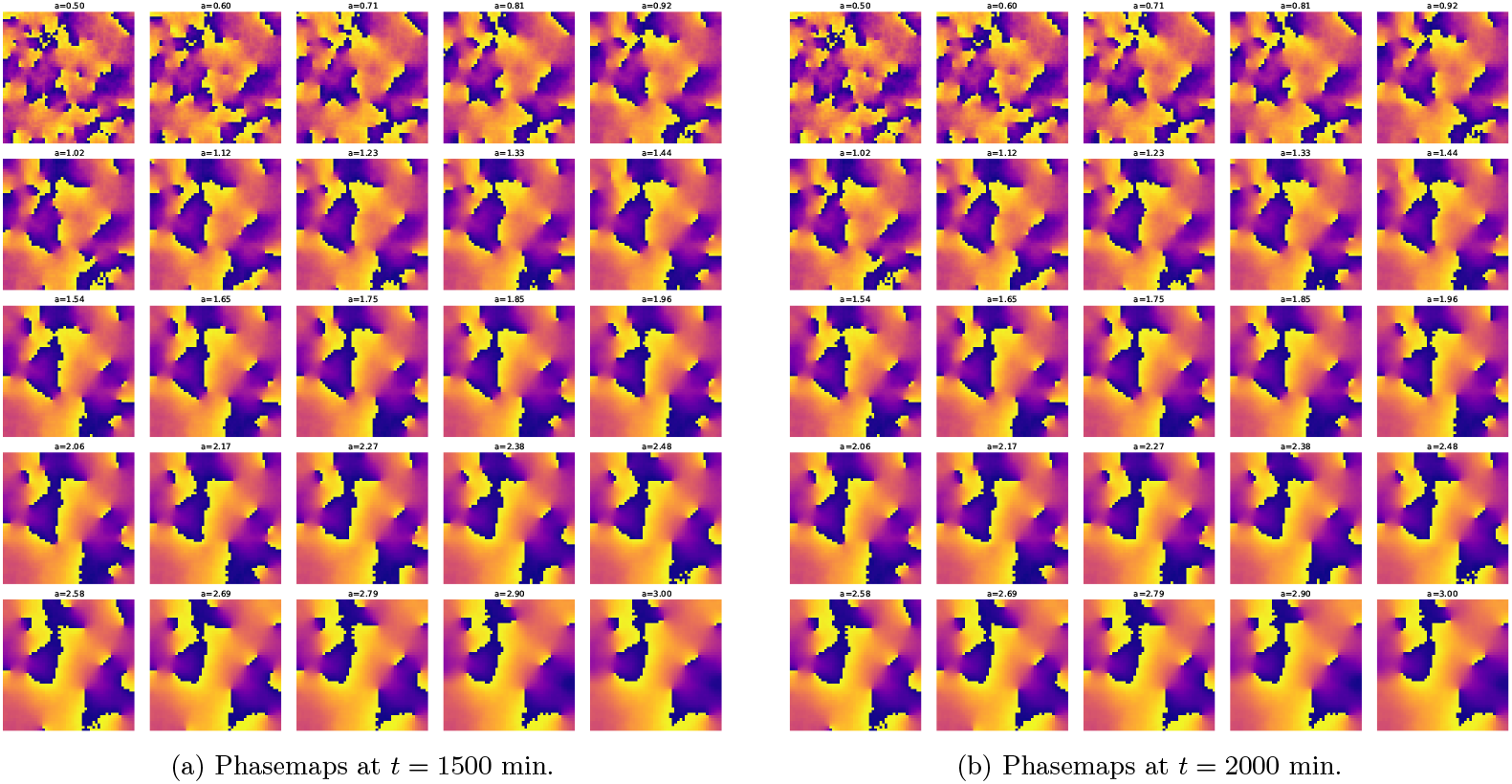
Phasemaps at *t* = 1500 and 2000 min obtained by simulating the 2D Kuramoto model with single sinusoidal coupling given by Eqs. (1). The initial phases are randomly sampled from a uniform distribution between [−*π, π*. At long times, we see topological defects known as vortices as previously reported in Refs. [6–8].

**FIG. 19:**
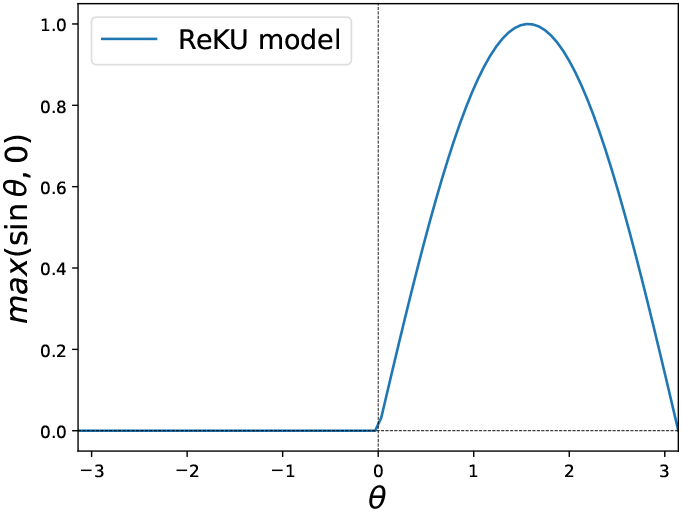
Coupling function for the ReKU model.

**FIG. 20:**
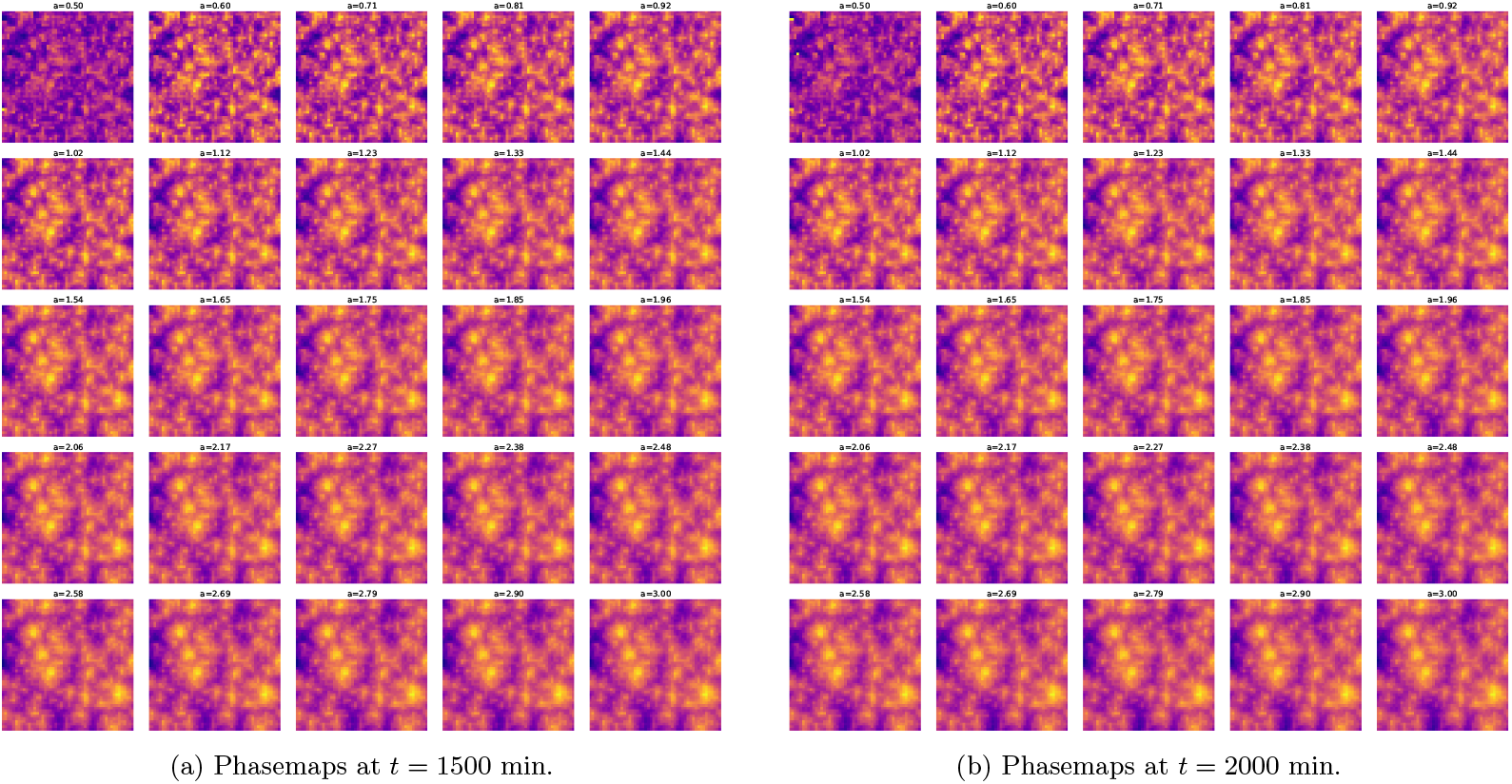
Phasemaps at *t* = 1500 and 2000 min obtained by simulating the 2D ReKU model described by Eqs. (3). The initial phases are randomly sampled from a uniform distribution between [ −*π/*2, *π/*2]. For the natural frequencies, we randomly sample them from a uniform distribution between [2*π/*180, 2*π/*150] min^−1^, since we would like to test these canonical models against experimental frequencies of PSM cells. 25 equally spaced values for the coupling constant *K* = *a*Δ_*ω*_ are used where *a* ∈ [0.5 −3]. Notice the phase homogeneity for the different coupling strengths at long times.

**FIG. 21:**
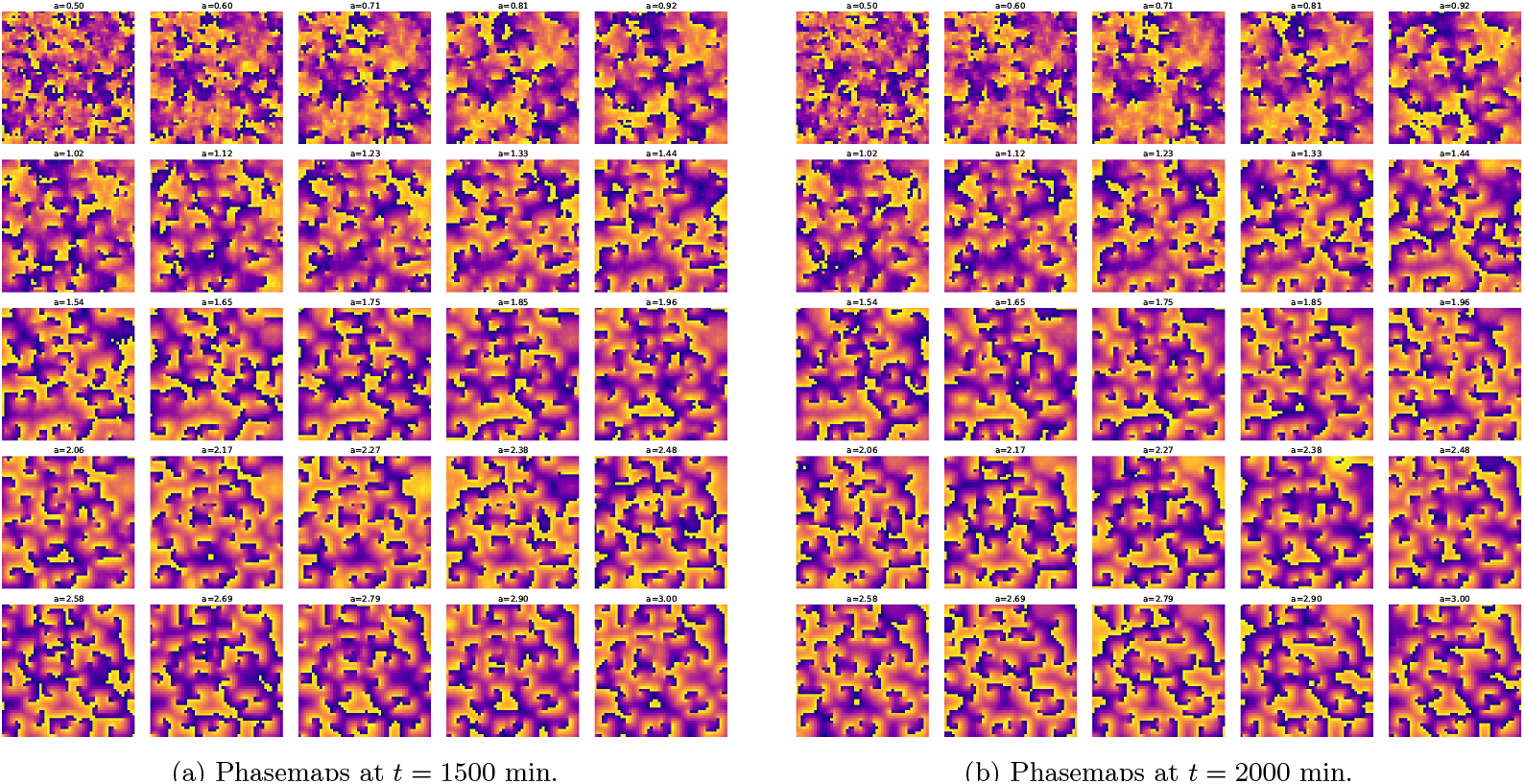
Phasemaps at *t* = 1500 and 2000 min obtained by simulating the 2D ReKU model. The initial phases are randomly sampled from a uniform distribution between [−*π, π*.

**FIG. 22:**
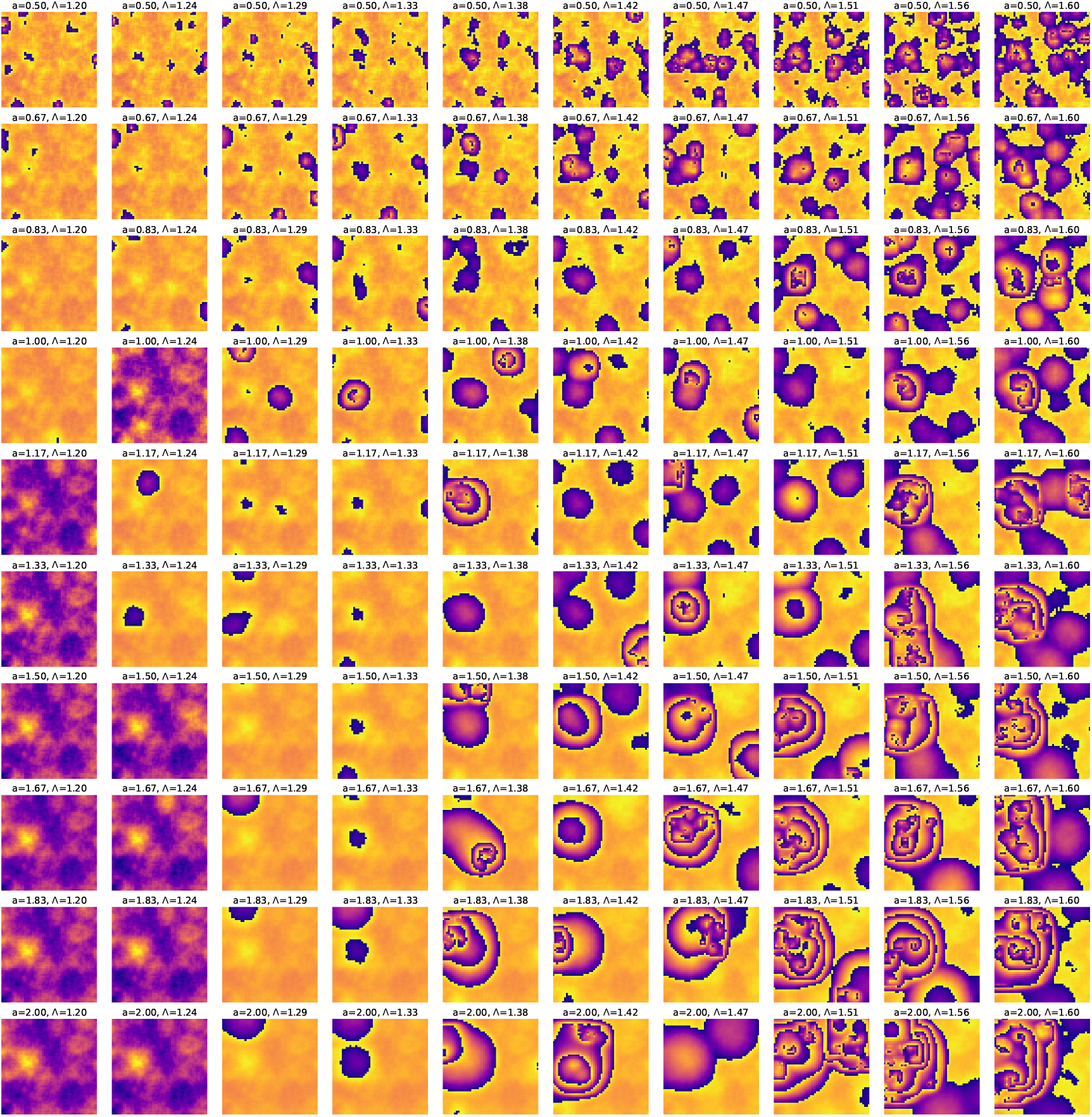
Phasemaps at *t* = 1000 min obtained by simulating the 2D Kuramoto model for QIF neurons described by Eqs. (4). The initial phases are randomly sampled from a uniform distribution between [ −*π/*2, *π/*2] and the natural frequencies are sampled randomly from a uniform distribution between [2*π/*180, 2*π/*150] min^−1^. The model parameters Λ and *K* = *a*Δ_*ω*_, where Δ_*ω*_ is the width of the frequency distribution and Λ are chosen from ten equally spaced values in Λ ∈ [1.2− 1.6] and *a* ∈ [0.5−2.0]. We do not observe concentric phase waves at multiple foci but only see quasiregular concentric phase wave patterns for certain (*K*, Λ) values indicating that the model cannot consistently generate such patterns for a wide range of fluctuating parameters.

**FIG. 23:**
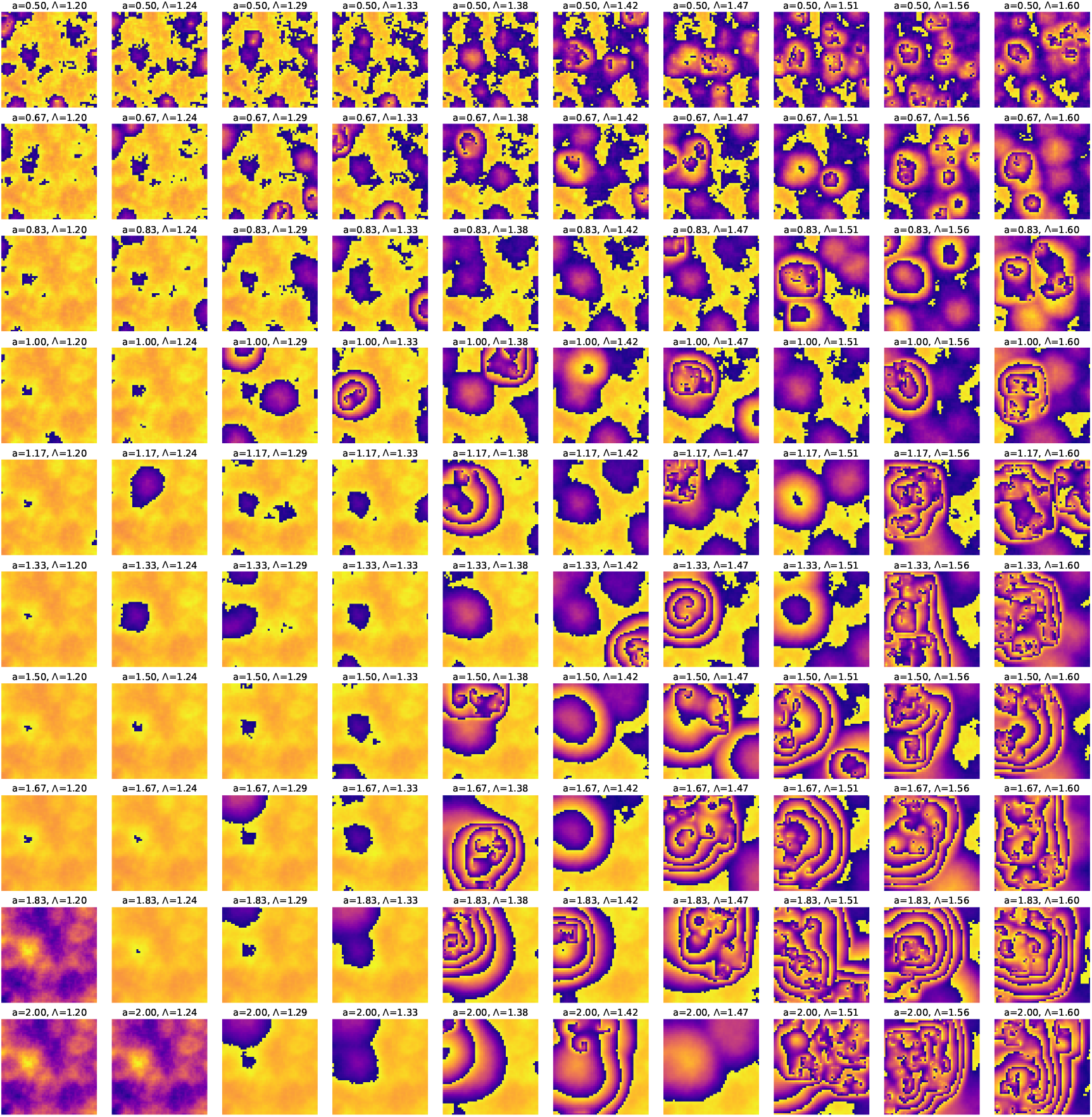
Phasemaps at *t* = 1500 min obtained by simulating the 2D Kuramoto model for QIF neurons described by Eqs. (4). The initial phases are randomly sampled from a uniform distribution between [ −*π/*2, *π/*2] and the natural frequencies are sampled randomly from a uniform distribution between [2*π/*180, 2*π/*150] min^−1^. The model parameters Λ and *K* = *a*Δ_*ω*_, where Δ_*ω*_ is the width of the frequency distribution and Λ are chosen from ten equally spaced values in Λ ∈ [1.2 − 1.6] and *a* ∈ [0.5 − 2.0].

**FIG. 24:**
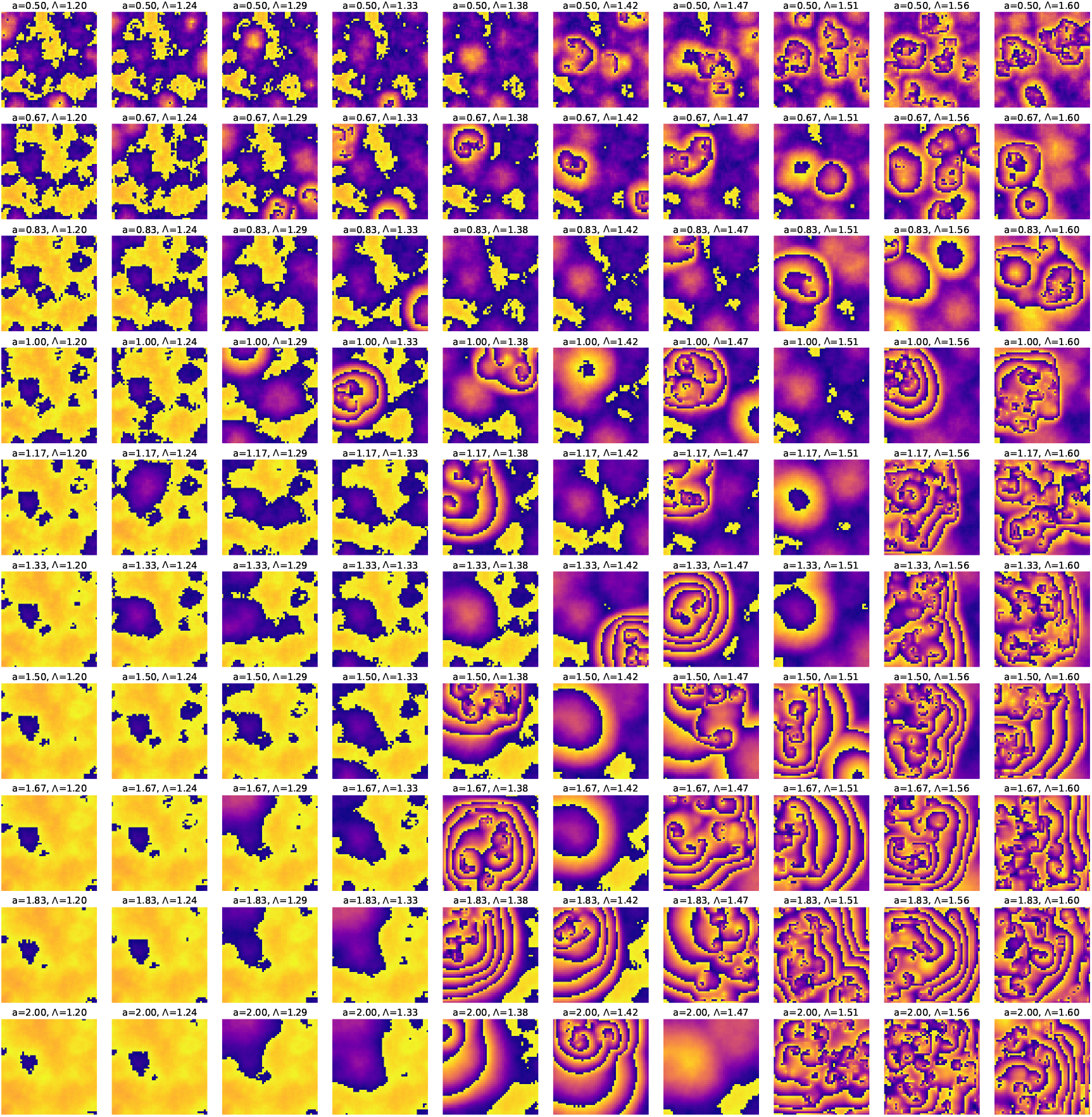
Phasemaps at *t* = 2000 min obtained by simulating the 2D Kuramoto model for QIF neurons described by Eqs. (4). The initial phases are randomly sampled from a uniform distribution between [ −*π/*2, *π/*2] and the natural frequencies are sampled randomly from a uniform distribution between [2*π/*180, 2*π/*150] min^−1^. The model parameters Λ and *K* = *a*Δ_*ω*_, where Δ_*ω*_ is the width of the frequency distribution and Λ are chosen from ten equally spaced values in Λ ∈ [1.2 − 1.6] and *a* ∈ [0.5 − 2.0]. Only a few target wave patterns survive at long times.

**FIG. 25:**
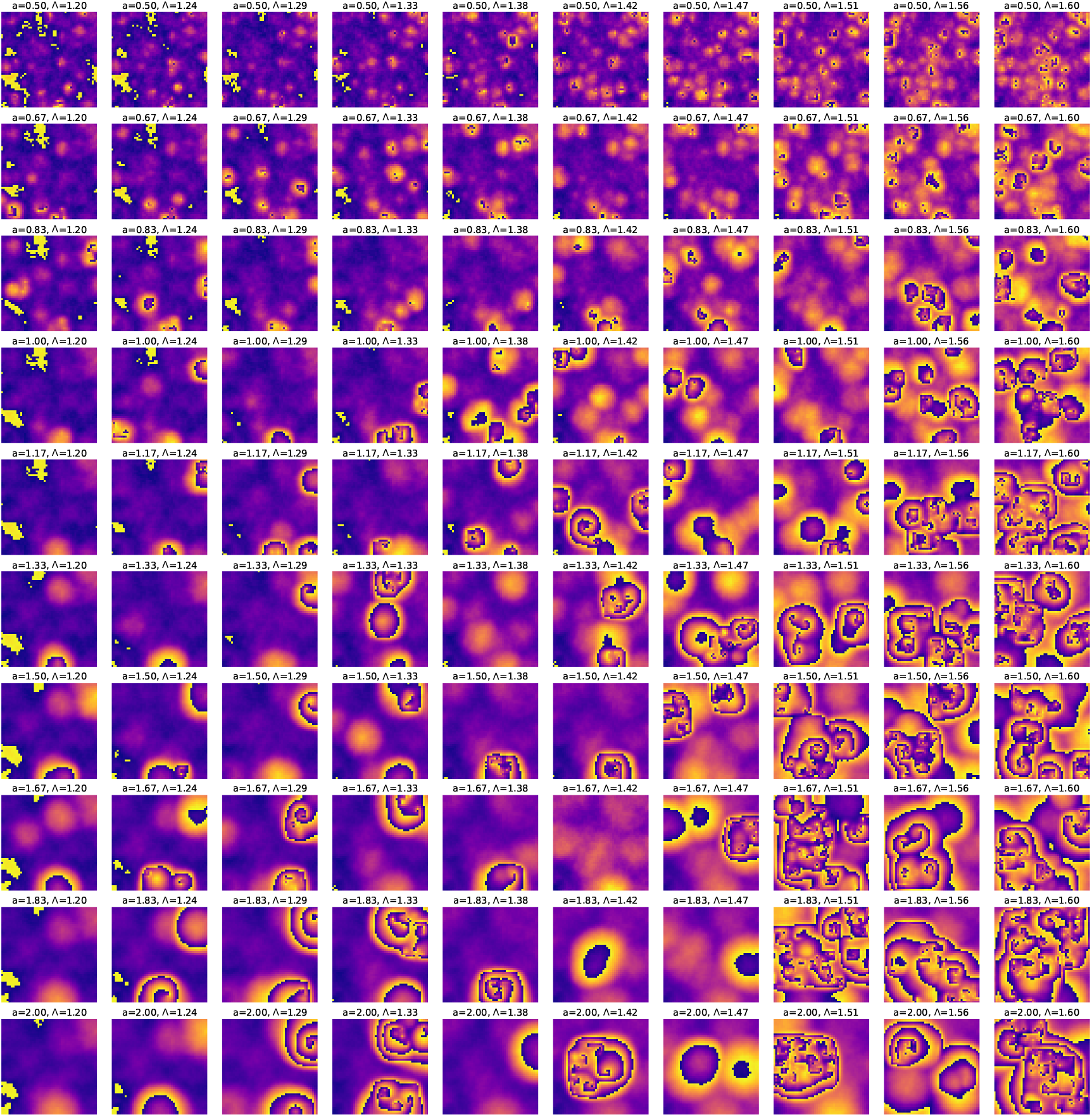
Phasemaps at *t* = 1000 min obtained by simulating the 2D Kuramoto model for QIF neurons described by Eqs. (4). The initial phases are randomly sampled from a much wider uniform distribution between [ −*π/*4, *π*] and the natural frequencies are sampled randomly from a uniform distribution between [2*π/*180, 2*π/*150] min^−1^. The model parameters Λ and *K* = *a*Δ_*ω*_, where Δ_*ω*_ is the width of the frequency distribution and Λ are chosen from ten equally spaced values in Λ ∈ [1.2 − 1.6] and *a* ∈ [0.5 − 2.0]. The subset of (*K*, Λ) values for which the model generates quasiregular target wave patterns is further limited compared to the initial phase distribution of width *π*. We see the emergence of spiral phase waves as the dominant temporal patterns.

**FIG. 26:**
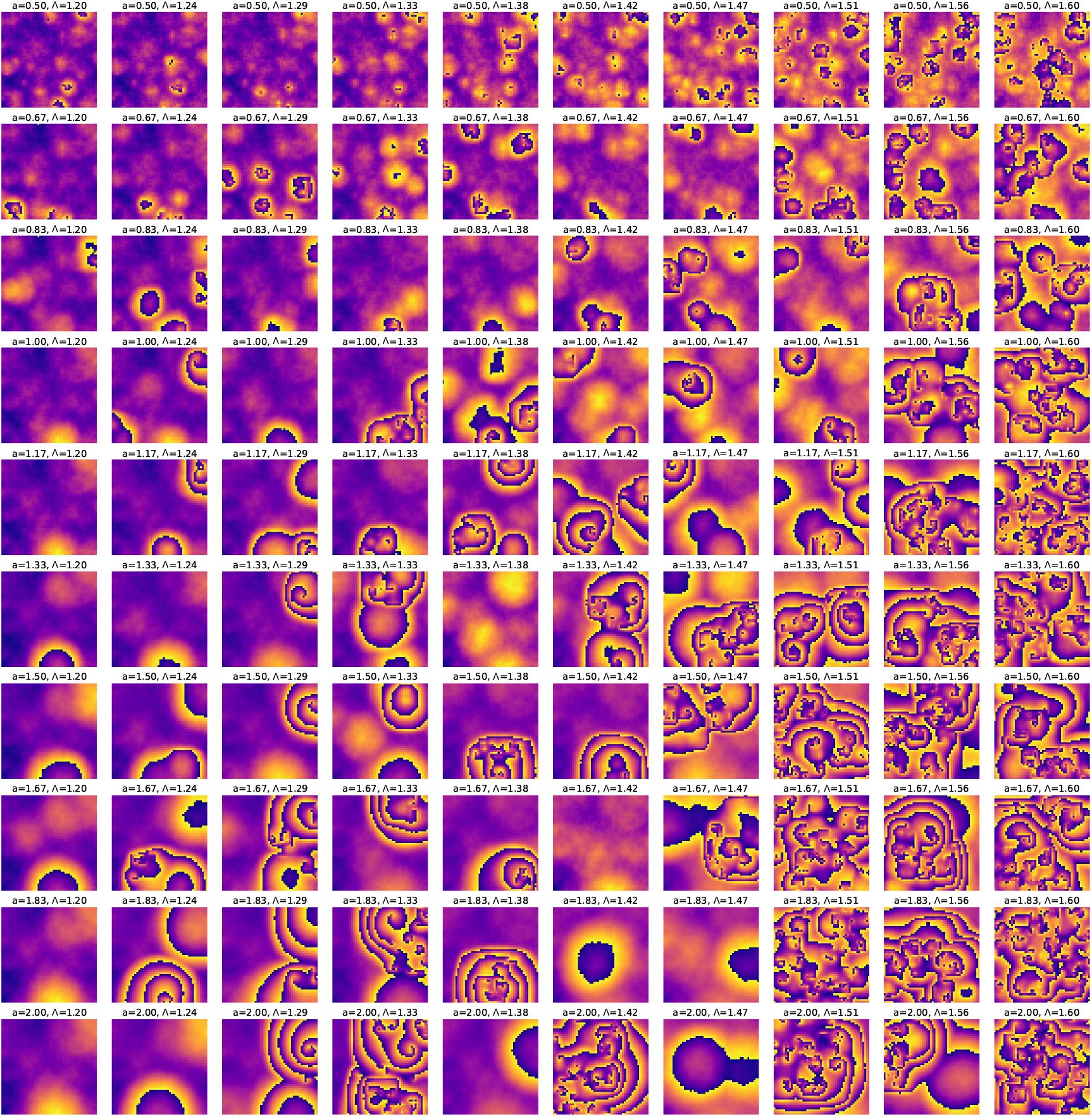
Phasemaps at *t* = 1500 min obtained by simulating the 2D Kuramoto model for QIF neurons described by Eqs. (4). The initial phases are randomly sampled from a much wider uniform distribution between [ −*π/*4, *π*] and the natural frequencies are sampled randomly from a uniform distribution between [2*π/*180, 2*π/*150] min^−1^. The model parameters Λ and *K* = *a*Δ_*ω*_, where Δ_*ω*_ is the width of the frequency distribution and Λ are chosen from ten equally spaced values in Λ ∈ [1.2 − 1.6] and *a* ∈ [0.5 − 2.0].

**FIG. 27:**
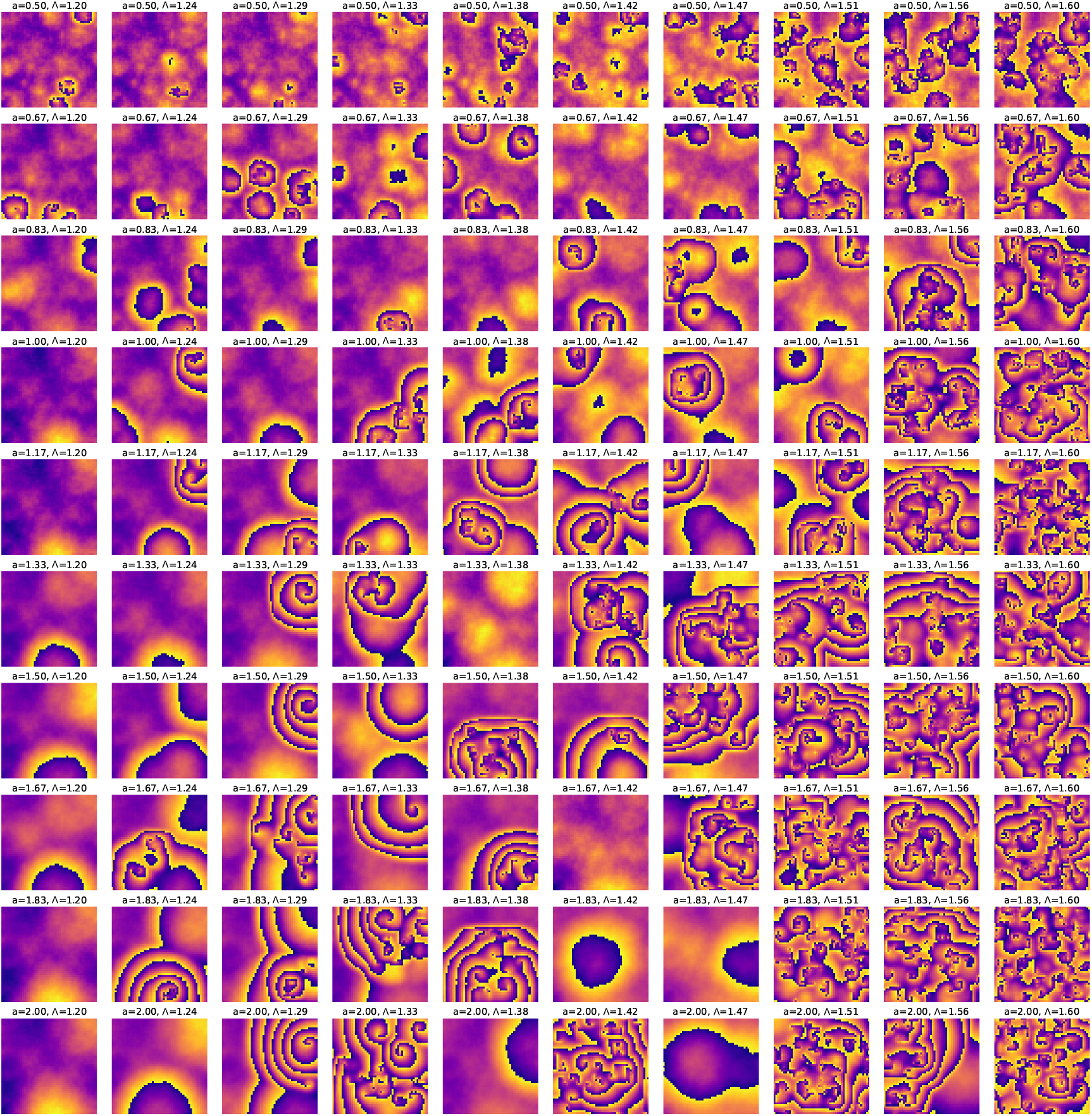
Phasemaps at *t* = 2000 min obtained by simulating the 2D Kuramoto model for QIF neurons described by Eqs. (4). The initial phases are randomly sampled from a much wider uniform distribution between [ −*π/*4, *π*] and the natural frequencies are sampled randomly from a uniform distribution between [2*π/*180, 2*π/*150] min^−1^. The model parameters Λ and *K* = *a*Δ_*ω*_, where Δ_*ω*_ is the width of the frequency distribution and Λ are chosen from ten equally spaced values in Λ ∈ [1.2 − 1.6] and *a* ∈ [0.5 − 2.0]. Very few parameter sets lead to sustained concentric phase wave patterns at long times.

